# Extensive remodelling of *XIST* regulatory networks during primate evolution

**DOI:** 10.1101/2023.12.04.569904

**Authors:** Emmanuel Cazottes, Charbel Alfeghaly, Cloé Rognard, Agnese Loda, Gaël Castel, Laura Villacorta, Michael Dong, Edith Heard, Irène Aksoy, Pierre Savatier, Céline Morey, Claire Rougeulle

## Abstract

Unravelling how gene regulatory networks are remodelled during evolution is crucial to understand how species adapt to environmental changes. We addressed this question for X-chromosome inactivation, a process essential to female development that is governed, in eutherians, by the *XIST* lncRNA and its *cis*-regulators. To reach high resolution, we studied closely related primate species, spanning 55 million years of evolution. We show that the *XIST* regulatory circuitry has diversified extensively over such evolutionary timeframe. The insertion of a HERVK transposon has reshuffled *XIST* 3D interaction network in macaque embryonic stem cells (ESC) and *XIST* expression is maintained by the additive effects of the *JPX* lncRNA gene and a macaque specific enhancer. In contrast, *JPX* is the main contributor to *XIST* expression in human ESCs but is not significantly involved in *XIST* regulation in marmoset ESCs. None of these entities are however under purifying selection, which suggests that neutrally evolving non-coding elements harbour high adaptive potentials.

## Introduction

Unravelling how phenotypic diversification is achieved across mammals is a long-standing question that becomes highly relevant in contexts of environmental variations. However, identifying and connecting molecular changes with functional and phenotypic outcomes is especially challenging since large-scale annotation projects have revealed that a major proportion of any mammalian genomes (∼80% of any given genome) is associated with a biochemical activity of some sort^1^. This includes nucleosome post-translational modifications, binding of transcription factors or biochemical machineries, which ensure general cellular functions but also participate in cell type specific gene regulation. In parallel, it has been estimated that only 5-10% of the human genome is under selective constraint^2,3^. This suggests that, outside of protein-coding sequences, most of the genome consists of stretches of nucleotides that evolve under either a neutral drift regimen or a fast-evolving rate, or that are specific of a given species. This also implies that biochemical activity is generally uncoupled from sequence conservation, which significantly hinders the prediction of the evolutionary potential of non-coding sequences.

Cis-Regulatory Elements (cRE) and long non-coding RNA genes (LRG) are thought to drive the evolution of gene regulatory networks. CREs, marked by specific features including chromatin accessibility, histone modifications such as H3K4me1 and H3K27ac, Transcription Factors (TF) binding or CTCF occupancy, constitute a significant proportion of the non-coding DNA^4^. They play crucial roles in spatiotemporal gene expression during development and act, at least in part, through physical interactions with target sequences (mostly enhancers) or participate in 3D genome partitioning (chromatin boundaries)^5–8^. On their side, long non-coding RNAs (LncRNA) are the most abundant transcriptional outcome of mammalian genomes and are generally lowly expressed, in a tissue-specific manner^9^. LRGs show intrinsic mechanistic plasticity since they may operate through the RNA molecule, the transcription process and/or the genomic locus itself^10^. Consequently, they exert *cis* and *trans*, transcriptional and post-transcriptional regulatory functions in a wide diversity of biological processes^11,12^. Many cREs and LRGs are however poorly conserved in sequence between species, which theoretically indicates a lack of function^13–16^. In mammals, a large fraction of these entities is randomly generated by the genetic drift that shapes the genome of multicellular eukaryotes^17^ and/or derives from Transposable Elements (TE), which provide platforms for TF binding and/or generate transcription start sites (TSS)^16,18–22^. Hence most cREs and LRGs emerge through non-adaptive neutral evolution and bring additional components to pre-existing regulatory loops^23–25^. While this increase in complexity may not immediately benefit to the species, it is thought to constitute an adaptive reservoir that can be recruited upon environmental changes and participate to phenotypic diversification^22,26^. Indeed cREs and LRGs have been correlated with the evolution of species-specific traits^15,16,27–31^. Yet, how functional redundancy is integrated during the evolution of a given regulatory system or how non-coding actors are re-hierarchized to orchestrate gene expression is still poorly studied.

X chromosome inactivation (XCI) provides an instrumental framework to interrogate the impact of variations in non-coding regulators on the evolution of gene regulatory networks. XCI is established during early female development and is mediated by the accumulation of *XIST* lncRNA on one of the two X-chromosomes, which triggers chromatin re-architecting including apposition of repressive histone marks H2AK119ub and H3K27me3 and transcriptional silencing of X-linked genes in *cis*^32,33^. *XIST* expression is a paradigm for epigenetic gene regulations during development: 1) *XIST* expression becomes monoallelic in specific lineages at specific developmental stages that vary across mammals and 2) *XIST* expression is then maintained throughout development with *XIST* RNA coating the inactive X (Xi) in most cells. These regulatory requirements are integrated by non-coding actors flanking the *XIST* gene within the X-inactivation centre (*XIC*). These include cREs^34–37^ and a compendium of LRGs^38–42^ the action of which is delimited by topological chromatin domains^43–45^. Significant differences have been reported in the way *XIST* is regulated by *XIC* actors between humans and mice, which may underly the divergence in timing and pattern of *XIST* expression during evolution^46–49^. In the mouse, *Xist* is under the control of repressive LRGs (*Tsix, Linx, Xite*) that have no functional orthologs in humans and of activator LRGs (*Jpx*, *Ftx* and *Xert*) that are either not operating in corresponding human cellular contexts or act through different *modus operandi*^36,41,50^. For instance, we have shown that *JPX* is a potent regulator of *XIST* in both human and mouse but acts through distinct modules (act of transcription and RNA respectively) to control the production of *XIST* at various steps depending on the species, while the function of *FTX* is not conserved in humans^41^.

To trace the chain of molecular events that shaped *XIST* regulation during evolution, we used unbiased functional approaches to identify putative players and test their role. To capture gradual or transient regulatory innovations, we focused on two closely related primate species, rhesus macaque and white tufted-eared marmoset that diverged from humans 35 and 55 million years ago, respectively. We first revealed a pronounced, TE-driven, 3D remodelling of the *XIST* interaction hub in rhesus embryonic stem cells (rhESCs) compared to humans. In parallel, we demonstrated that *JPX* regulatory activity toward *XIST* is strongly reduced in macaques compared to humans and only controls *XIST* expression levels and not the monoallelic accumulation of *XIST* RNA on the Xi. In addition, *JPX* acts in combination with a macaque specific enhancer to maintain *XIST* expression levels in rhESCs. We further showed that *JPX* does not significantly regulate *XIST* in female marmoset embryonic stem cells (cjESCs), suggesting a gradual functionalization of *JPX* LRG in primate species. Since sequence comparison of these non-coding players did not show evidence of purifying selection, our observations suggest that non-adaptive evolution plays a major role in the remodelling of the *XIST* regulatory network in primates. Altogether, our results illustrate how combined evolutions of LRGs, TE-mediated 3D reorganisation and cRE concur to the evolution of regulatory networks across short evolutionary timescales.

## RESULTS

### Rhesus pluripotent stem cells have established X-inactivation

To study the regulation of *XIST* in non-human primates, we used a female rhesus macaque embryonic stem cell (rhESC) line (LYON-ES1) which models early developmental stages^51^. RNA-seq analyses of LYON-ES1 indicate a transcriptional signature closely resembling that of the post-implantation epiblast (day 13–14 post-fertilization) of the macaque embryo (Supplementary Fig. 1A). As such, rhESCs represent the macaque equivalent of primed human embryonic stem cells (hESCs)^52,53^.

To determine the XCI status in LYON-ES1 rhESCs, we first performed RNA-FISH for *XIST*. The vast majority of nuclei (92%) exhibited one domain of *XIST* accumulation (Fig. 1A). Immuno-RNA-FISH further showed a co-localisation of *XIST* domains with foci of H3K27me3 (Fig. 1B) and of H2AK119Ub (Fig. 1C) in 91% or 95% of nuclei respectively, which indicates a co-enrichment of these repressive histone marks on the *XIST*-coated X chromosome. To assess the transcriptional activity of X-linked genes, we then performed RNA-FISH for *XIST* and for two X-linked genes, *ATRX* and *POLA1*, located on the long and short arms of the macaque X chromosome respectively (Fig. 1D). For both genes, a single transcription pinpoint was detected away from the *XIST* domain indicating mono-allelic transcription from the presumptive active X (Xa) and silencing of alleles on the *XIST*-coated X (Fig. 1D). Consistently, allelic analyses of RNA-seq data of LYON-ES1^54^ showed an allelic ratio of 0.2 for informative X-linked positions (i.e., reads overlapping a SNP, see Methods), indicating preferential expression of one haplotype. This contrasts with allelic ratios at genes on chromosome 7 or 8, which were centred on 0.5 which, reflects bi-allelic expression (Fig. 1E).

**Figure 1:**
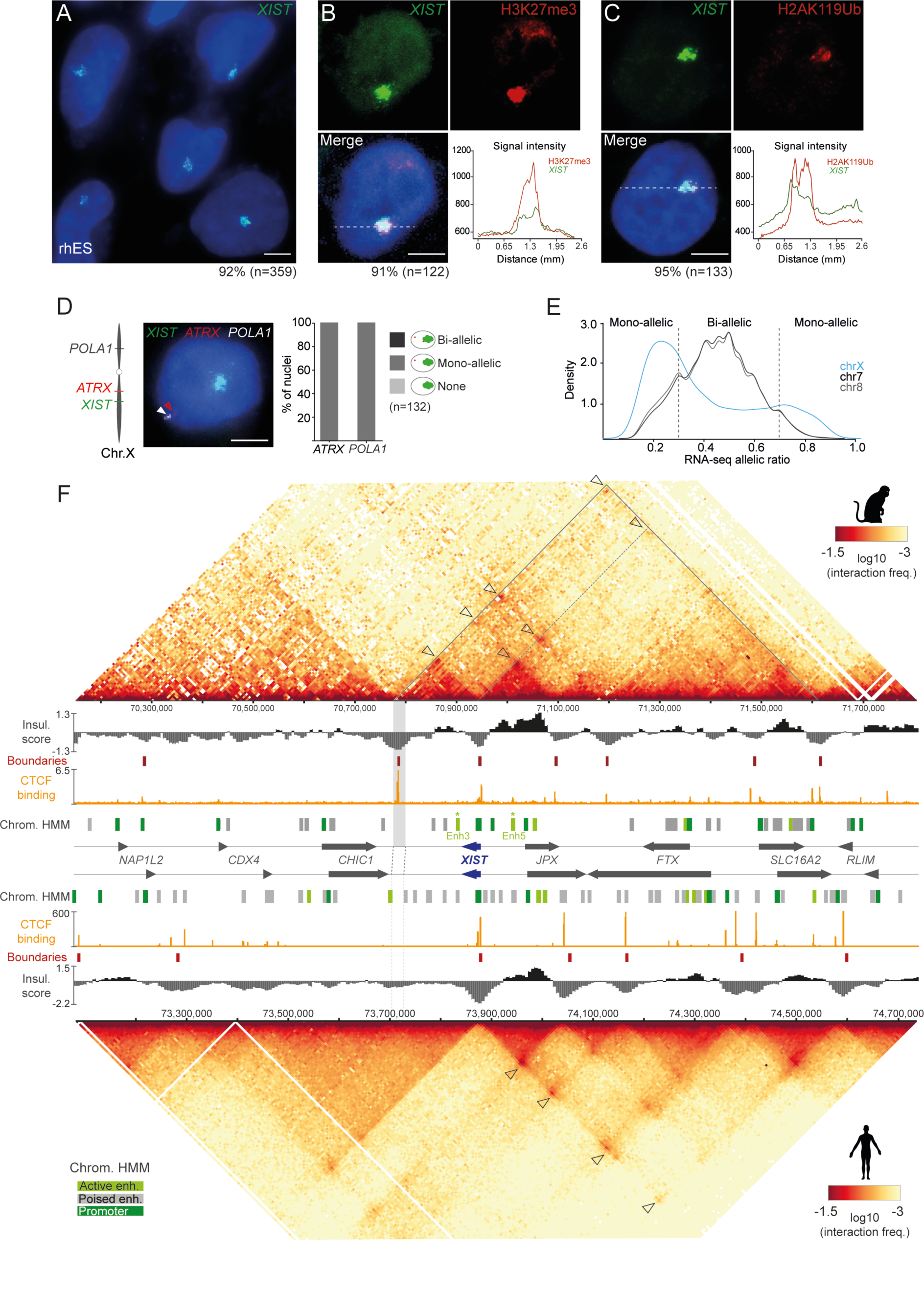
X-inactivation in rhESCs is associated with a specific chromatin landscape of the *XIC*. A. Representative image of *XIST* RNA-FISH in LYON-ES1 rhESCs. The percentage of nuclei with an *XIST* RNA accumulation is indicated. (Scale bar: 5 µm) B. Combined *XIST* RNA-FISH (green) and H3K27me3 immunofluorescence staining (red) in LYON-ES1 rhESCs. The number of nuclei with a *XIST* domain is indicated. The distributions of each signal intensity along the depicted dotted line are shown for one representative nucleus. (Scale bar: 5 µm) C. Same as panel B for H2AK119Ub immunofluorescence staining. D. Triple RNA-FISH for *XIST* (green), *ATRX* (red) and *POLA1* (white) in LYON-ES1 rhESCs. Gene positions on the macaque X chromosome are indicated. The bar plot shows the percentages of indicated expression patterns in the cell population. (Scale bar: 5 µm) E. Density plots of allelic ratios at polymorphic positions on chromosome X, 7 or 8 extracted from LYON-ES1 rhESCs RNA-Seq data^54^ (n=2). F. Capture-HiC maps of the *XIC* in LYON-ES1 rhESCs (top) and in H9 hESCs (bottom) showing interaction frequencies calculated over 6-kb bins (n=2). The distributions of insulation scores, the positions of boundaries between chromatin domains and CTCF binding profiles determined by CUT&RUN in LYON-ES1 rhESCs (n=2; this study) or by ChIP-seq in H9 hESCs (ENCODE) are shown. CTCF-anchored chromatin loops are pointed with arrowheads. The macaque-specific chromatin boundary is highlighted in light grey. The ChromHMM tracks depict the position of chromatin signatures marking active enhancers (light green), poised enhancers (grey) and promoters (dark green) in LYON-ES1 rhESCs and in H9 hESCs computed from CUT&RUN data and DNase I-seq data (n=2; this study and ENCFF291ZMA) (see Methods). Putative enhancers (Enh3 and Enh5) active in rhESCs specifically are indicated.

We concluded from these observations that the LYON-ES1 rhESC line is in a primed state of pluripotency with established XCI. Analyses performed on an independent rhiPSC line^55^ gave similar results (Supplementary Fig. 1B-D).

### Rhesus and human ESCs exhibit distinct *XIST* regulatory landscapes

To probe for *XIST* cREs in an unbiased manner and to directly compare *XIST cis*-regulatory networks in rhesus macaques and humans, we generated high-resolution chromatin contact maps within the *XIC* of female LYON-ES1 rhESCs and H9 hESCs. To this end, we performed capture HiC (cHiC)^56^ with tailored oligo probe sets covering a region of 3 Mb centred on the rhesus and on the human *XIST* genes (Supplementary Fig. 2A). Of note, probe design did not allow the exact same level of coverage of rhesus and human *XIC*s resulting in a slightly reduced resolution of the macaque interaction map. Since, for each species, biological replicates were highly correlated (r2 = 0.8 for hESCs and r2 = 0.60 for rhESCs) (Supplementary Fig. 2B), the interaction frequencies of the two replicates have been aggregated in the maps shown in Fig. 1F.

General inspection of macaque and human contact maps first revealed topological partitioning of the *XIC* in two domains in both species. A large centromeric domain covers the protein-coding genes *NAP1L2*, *CDX4* and *CHIC1*, and a more structured region encompasses the *XIST* non-coding gene and its known positive regulators *JPX* and *FTX* (Fig. 1F). In humans, the latter consists of smaller domains, with chromatin loops bridging the *XIST* promoter to specific loci along the *JPX-FTX* transcription units. These include interaction hotspots that have been recently proposed to contribute to *JPX* regulatory function^41^, but also elements in the intergenic region between *SLC16A2* and *RLIM*.

More detailed analysis revealed that a macaque-specific boundary located downstream of *CHIC1* has hijacked some of the interactions involving the *XIST* promoter (Fig. 1F). This boundary is characterized by the lowest insulation score of the sampled region, indicative of the strongest barrier activity of the locus. CTCF binding, profiled by CUT&RUN, shows that chromatin loops and interaction domains are anchored or segregated by CTCF binding in both species, as previously described at other loci^57–59^; this analysis further revealed that the macaque-specific boundary exhibits the highest CTCF binding signal of the rhesus *XIC* (Fig. 1F).

To identify candidate cREs, we compared cHiC data to CUT&RUN data for histone modifications marking promoters (H3K4me3)^60^ and enhancers (H3K27ac and H3K4me1) (this study), as well as chromatin accessibility maps in LYON-ES1 rhESCs (ATAC-seq, this study) and in hESCs (DNase I sensitivity; ENCODE) (Supplementary Fig. 2C and 2D). To ease functional interpretation of the results, we used ChromHMM^61^ to segregate the sequencing data into promoter-like, active enhancer-like and poised enhancer-like signatures (see Methods). While promoter signatures were generally highly conserved in both species, active enhancers appeared more variable between rhESCs and hESCs, in agreement with other genome-wide reports^2,15,16,31^ (Fig. 1F). In particular, we identified two putative active enhancers, namely enh3 and enh5, within the *XIST* topological domain in rhESCs, which are not detected in hESCs (Fig. 1F).

Altogether, these analyses reveal significant differences in the chromatin landscape of the *XIC* between rhESCs and hESCs. The interaction network involving the macaque *XIST* promoter is linked to a macaque-specific domain boundary downstream of *XIST* and to the presence of two candidate cREs in *XIST* promoter vicinity that are not active in hESCs. These reshaping of *XIST* contact maps suggest that distinct regulatory modes may be at work in these two primate species.

### Insertion of a HERVK element creates a rhesus-specific XIC topology

The macaque-specific chromatin boundary downstream of *XIST* consists of a TE of the Human Endogenous Retrovirus K (HERVK) family, which shelters four CTCF binding sites known to promote barrier activities^59^ (Fig. 2A, Supplementary Fig 3B). To trace the evolutionary history of the HERVK insertion, we blasted its sequence against the genomes of representative species of the primate lineage. No HERVK element could be detected in the new-world monkey (Platyrrhini) or pro-simian lineages, indicating that HERVK TEs invaded the genome of the old-world monkeys’ (Cercopithecus) and great apes’ (Hominoids) common ancestor after the divergence from the new-world monkeys approximately 40 to 25.5 Mya (Fig. 2B). In contrast, when we compared *XIC* sequences of selected primate species, we only found syntenic HERVK insertion in Cercopithecus species, which dates the HERVK insertion into the macaque *XIC* after the divergence with great-apes, from 25.5 to 16 Mya (Fig. 2B).

**Figure 2:**
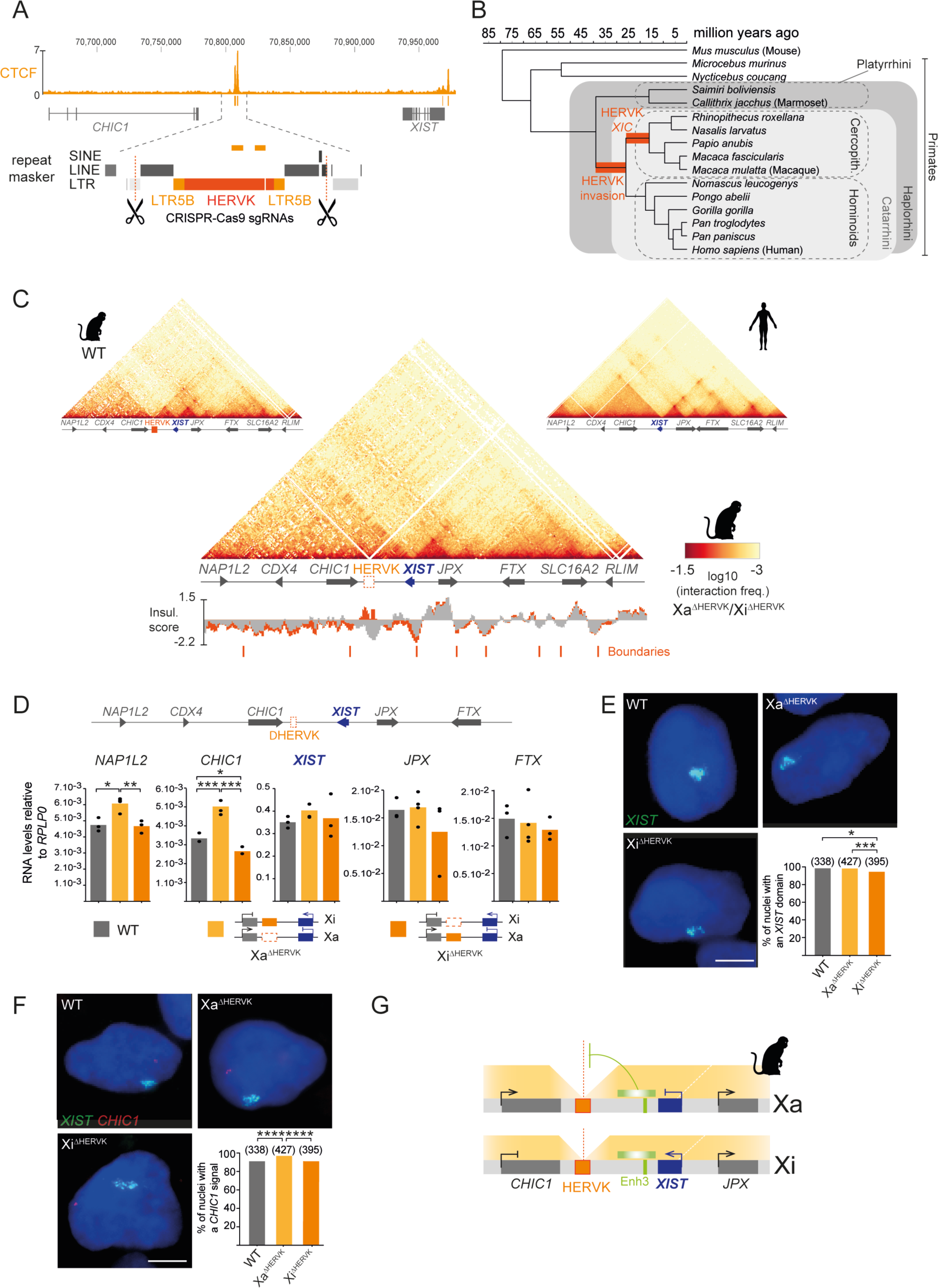
Humanizing the *XIC* chromatin architecture in rhESCs perturbs *NAP1L2* and *CHIC1* gene expression on the Xa. A. Schematic representation of the HERVK deletion strategy showing the position of the two guide RNAs with respect to surrounding repeated sequences (RheMac10 RepeatMasker annotation) and to CTCF CUT&RUN signals. B. Phylogenetic tree of representative primate species used to date HERVK insertion within the *XIC*. Branch lengths represent the time since divergence in Millions of years. C. Capture HiC contact map of the *XIC* in LYON-ES1 rhESCs carrying HERVK homozygote deletion compared to WT LYON-ES1 rhESCs (top left) or to WT H9 hESCs (top right). The superimposed distributions of insulation scores in WT (grey) and in deleted cells (orange) are shown. D. Average expression levels of indicated *XIC* genes measured by RT-qPCR in WT rhESCs, and in clones in which the HERVK is deleted on the Xa (Xa^ΔHERVK^) or on the Xi (Xi^ΔHERVK^). Dots represent the expression levels in independent clones. Unpaired two-sided t-test: *p-value < 0.05; **p-value < 0.01; ***p-value < 0.001. E. Representative images of *XIST* RNA-FISH in WT, Xa^ΔHERVK^ and Xi^ΔHERVK^ rhESCs. The percentages of cells with a *XIST* cloud are shown for each genotype on the bar plot. Fisher test: *p-value < 0.05; ***p-value < 0.001. Scale bar = 5 µm. F. Representative images of double RNA-FISH for *XIST* and for *CHIC1* in WT, Xa^ΔHERVK^ and Xi^ΔHERVK^ rhESCs. The percentages of cells with a *CHIC1* transcription pinpoint are shown for each genotype on the bar plot. Fisher test: ****p-value < 0.0001. Scale bar = 5 µm. G. Proposed model of HERVK function: a macaque specific boundary (dark orange) shields *CHIC1* and *NAP1L2* from ectopic interactions with a putative enhancer (Enh3, green) located between the HERVK and the *XIST* locus. The dotted white line shows the relocation of the boundary upon HERVK deletion.

Despite being annotated as an intact HERVK element by RepeatMasker, the sequence of the boundary has diverged from the HERVK consensus; notably ORFs are degenerated (data not shown). Among the four CTCF binding sites (CBS) embedded in the HERVK, CBS1, CBS2 and CBS4 are conserved across Cercopithecus species and CBS3 is gained in both *Macaca fascicularis* and *Macaca mulatta* (Supplementary Fig. 3A, B). The accumulation of CTCF binding sites is likely to provide higher insulating strength over CTCF binding at the *XIST* promoter, thereby creating an extra topological domain boundary in macaques.

### HERVK-associated boundary shields centromeric genes from *XIST* regulatory environment

To probe the function of the HERVK-mediated boundary in the 3D organization of the rhesus *XIC* and in the regulation *XIC* genes, we engineered a CRISPR/Cas9 deletion targeting the whole TE (Fig. 2A and Supplementary Fig. 4A). We selected independent LYON-ES1 rhES clones carrying the deletion on the Xa (Xa^ΔHERVK^), on the Xi (Xi^ΔHERVK^) or on both alleles as determined using a SNP within the deleted interval (Supplementary Fig. 4B). The deletion had no detectable effect on cell viability, cell renewal or pluripotency markers expression (Supplementary Fig. 4C). CHiC analysis showed a complete loss of barrier activity at the site of HERVK homozygote deletion, associated with strengthened insulation at the *XIST* promoter boundary (∼1.7 fold; Fig. 2C). Hence, this mutation recapitulates, in rhESCs, the *XIC* chromatin conformation of H9 hESCs. This demonstrates the impact of this HERVK element in shaping the topological organization of the rhesus *XIC*.

We then took advantage of Xa^ΔHERVK^ and Xi^ΔHERVK^ allelic configurations to assess the consequences of humanized *XIC* architecture on local gene expression in *cis*: on genes expressed from the Xa (*NAP1L2, CHIC1, JPX, FTX, SLC16A2* and *RLIM*) or from the Xi (*XIST* and *JPX*). The deletion had no significant effect on the expression of genes on the telomeric side of the HERVK boundary (*XIST*, *JPX*, *FTX*, *SLC16A2* and *RLIM*) in either allelic configuration (Fig. 2D and Supplementary Fig. 4D). Consistently, in ΔHERVK rhESCs, more than 90% of nuclei displayed a *XIST* domain as in WT cells, even though Xi^τιHERVK^ cells showed significantly less accumulations than their Xa^τιHERVK^ and WT counterparts (Fig. 2E). In contrast, we detected a significant increase in *CHIC1* and *NAP1L2* RNA levels in Xa^τιHERVK^ rhESCs compared to WT or to Xi^τιHERVK^ cells (Fig. 2D), associated with a higher percentage of nuclei exhibiting a *CHIC1* transcriptional pinpoint away from the *XIST* coated X by RNA-FISH (Fig. 2F).

These data suggest that the insulation activity of the HERVK TE does not impact *XIST* expression but instead protects, on the Xa, the centromeric genes *CHIC1* and *NAP1L2* from being contacted by putative regulatory sequences located in the *XIST* domain (Fig. 2G). Consistently, analyses of cHiC data using *CHIC1* or *NAP1L2* promoters as viewpoints revealed increased interaction frequencies with few discrete regions downstream of *CHIC1* including Enh3 (Supplementary Fig. 4E). In addition, the insulation activity at the *XIST* promoter is significantly increased upon HERVK deletion (Fig. 2C). Of note, the Xa-restricted effect of the deletion indicates that *CHIC1* and *NAP1L2* promoters on the Xi are not sensitive to regulatory influences originating from the *XIST* domain.

### *JPX* is a minor regulator of *XIST* in rhesus macaque ESCs

The rhesus *JPX-FTX* locus harbours active enhancer signatures and is engaged in chromatin loops with the *XIST* promoter (Fig. 1F). In addition, *JPX* is a known positive regulator of *XIST* in mice and humans^41,62^, while *FTX* exerts such a function in mice only^40,41^. During female macaque pre-implantation development, *XIST*, *JPX* and *FTX* are concomitantly up-regulated (Fig. 3A) and their expression dynamics are strongly correlated compared to other *XIC* genes (Fig. 3B). This suggests that *JPX* and *FTX* could participate in *XIST* regulation in macaques. RNA-FISH analysis of *JPX* and *FTX* transcription in rhESCs showed that *JPX*, but not *FTX*, escape XCI, indicating that, like in humans, *JPX* could exert a *cis*-regulatory function (Fig. 3C). Rhesus *JPX* however is transcribed from the Xi in only ∼50% of LYON-ES1 rhESCs while human *JPX* escapes XCI in nearly all cells of the H9 hES cell line^41^.

**Figure 3:**
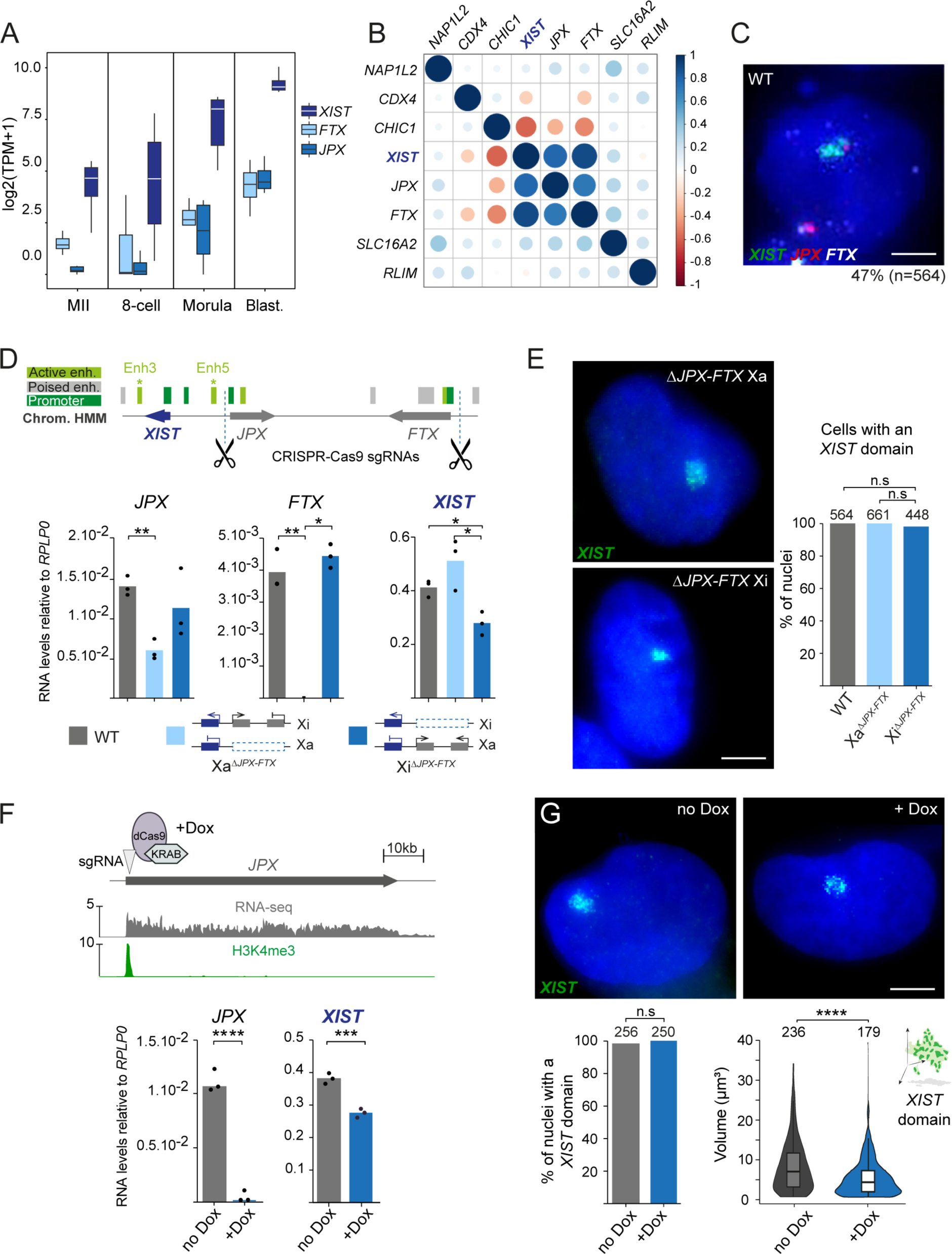
*JPX* inhibition, like τι*JPX-FTX,* weakly impacts *XIST* expression in rhESCs. A. *XIST*, *JPX* and *FTX* expression dynamics during pre-implantation development of female rhesus macaque extracted from RNA-seq dataset GSE112534^54^. B. Pearson correlation between *XIC* gene expression profiles during pre-implantation development of female rhesus macaques computed from RNA-seq dataset GSE112534^54^. C. Representative picture of triple RNA-FISH for *XIST*, *JPX* and *FTX* in LYON-ES1 rhESCs. The percentage of cells with biallelic transcription of *JPX* associated with monoallelic expression of *FTX* is indicated. D. CRISPR-Cas9 deletion strategy targeting the *JPX-FTX* genomic interval showing the position of the guide RNA relative to the genes and to chromatin features. Average expression levels of *JPX*, *FTX* and *XIST* measured by RT-qPCR in WT cells and in cells in which the *JPX-FTX* locus is deleted on the Xa (Xa^ΔJPX-FTX^) or on the Xi (Xi^ΔJPX-FTX^) (n=3). Dots represent expression levels in independent clones. Unpaired two-sided t-test: *p-value < 0.05; **p-value < 0.01. E. Representative pictures of *XIST* RNA-FISH in LYON-ES1 rhESCs carrying the *FTX-JPX* deletion on the Xa or on the Xi. The percentages of cells with a *XIST* cloud are shown on the bar-plot aside. Fisher test: n.s., not significant. Scale bar = 5 µm. F. Dox-inducible CRISPRi strategy showing the position of the guide RNA targeting the *JPX* promoter, RNA-Seq profile and H3K4me3 CUT&RUN distribution along the *JPX* transcription unit of LYON-ES1 rhESCs. Below, average expression levels of *JPX* and *XIST* measured by RT-qPCR in *JPX* CRISPRi rhESCs treated (+Dox) or not (no Dox) with doxycycline (n=3). Unpaired two-sided t-test: ***p-value < 0.001; ****p-value < 0.0001. G. Representative images of *XIST* RNA-FISH in *JPX* CRISPRi rhESCs treated (+Dox) or not (no Dox) with doxycycline. The percentages of cells with a *XIST* cloud are shown on the bar-plot underneath. Fisher test: n.s., not significant. Violin plots show the distributions of the volumes of *XIST* RNA accumulations. Wilcoxon test: ****p-value < 0.0001. Scale bar = 5 µm.

To test the regulatory potential of the *JPX-FTX* region, we generated independent LYON-ES1 rhESC clones bearing a large (∼330 kb) deletion on either allele (Fig. 3D; Supplementary Fig. 5A and 5B). Mutant cells did not display overt phenotype and expressed normal levels of markers of ESC identity (Supplementary Fig. 5C). As expected, *FTX* expression was completely abrogated in Xa^1ι*JXP-FTX*^ clones, while *JPX* RNA levels were significantly reduced (Fig. 3D), due to remaining expression from the Xi, as confirmed by RNA-FISH (Supplementary Fig. 5D). No significant effect on *XIST* expression could be detected either by RT-qPCR (Fig. 3D) or by RNA-FISH in this allelic configuration (Fig. 3E). In cells carrying the deletion on the Xi, *FTX* allelic transcription pattern and *FTX* RNA levels appeared unchanged compared to WT cells, while *JPX* transcription from the Xi was lost and transcript abundance was reduced with some variability between the clones (Fig. 3D and Supplementary Fig. 5D). In this genotype, we observed a mild but significant reduction of *XIST* RNA levels compared to WT (∼1.6-fold) or to Xa^1ι*JXP-FTX*^ (∼2-fold) cells (Fig. 3D). This was confirmed by RNA-FISH analyses, which revealed a slight reduction in the volume of the *XIST* domain on the Xi^1ι*JXP-FTX*^ (Supplementary Fig. 5E). We, however, did not detect any significant change in the proportion of cells displaying a *XIST* accumulation (Fig. 3E), and the silencing of X-linked genes *POLA1* and *ATRX* was not affected (Supplementary Fig. 5F). Thus, the *JPX-FTX* locus exerts a minor *cis-*regulatory activity on *XIST* expression. Since no other *XIC* genes were affected by the mutation, irrespective of the deletion allelism, we concluded this mild *cis-*regulatory activity targets *XIST* specifically.

The deletion of the entire *JPX-FTX* locus in rhESCs allows to address the regulatory potential of a locus containing several candidates but may mask the function of specific elements. We thus interrogated more specifically the function of *JPX* since it is expressed from the Xi and has a preponderant role in *XIST* regulation in hESCs^41^. To this end, we knocked down *JPX* using a doxycycline-inducible CRISPRi system featuring a catalytically dead Cas9 protein fused to a KRAB repressor domain^63,64^ (Fig. 3F). We selected independent LYON-ES1 clones with a minimal number of transgenic insertions, as determined by qPCR on genomic DNA, to minimize artefactual effects induced by the random insertion of the transgenes (Supplementary Fig. 5G). Upon doxycycline induction of the dCas9-KRAB, *JPX* was efficiently repressed as measured by RT-qPCR and RNA-FISH (Fig. 3F and Supplementary Fig. 5H). *XIST* RNA levels appeared mildly diminished (∼1.3-fold) following *JPX* KD compared to non-induced cells (Fig. 3F) and we detected *XIST* RNA accumulations – although smaller in volume – in most cells, similarly to what we observed in Xi^τι*JXP-FTX*^ rhESCs (Fig. 3G).

Altogether, these results show that, like in mice and humans, the *JPX-FTX* locus regulates *XIST* in *cis* in rhESCs. We could attribute with high confidence this function to *JPX*, since *JPX* KD closely reproduces the phenotype of the deletion of the *JPX-FTX* span. Yet, *JPX* exerts a much weaker regulatory activity in macaque ESCs compared to human and mice and appears to subtly control *XIST* expression levels rather than the probability of *XIST* to be activated, repressed or to coat the Xi.

### *JPX* does not significantly participate in *XIST* regulation in white-tufted-ear marmoset ESCs

This unexpected result prompted us to track the evolution of *JPX* as an *XIST* regulator in primates. We therefore turned to the white-tufted-ear marmoset (*Callithrix jacchus)*, a representative of the new-world monkey clade, which diverged 39 Mya from the ancestor that gave rise to the old-world monkeys and great apes (Fig. 2B). This species should be instrumental in studying the ancestral state of the regulatory relationship between *JPX* and *XIST* in primates. During female marmoset pre-implantation development, *XIST*, *JPX*, and *FTX* are co-expressed (Fig. 4A), but their expression is not significantly correlated with that of *XIST*, as opposed to the situation in the macaque (Fig. 4B).

**Figure 4:**
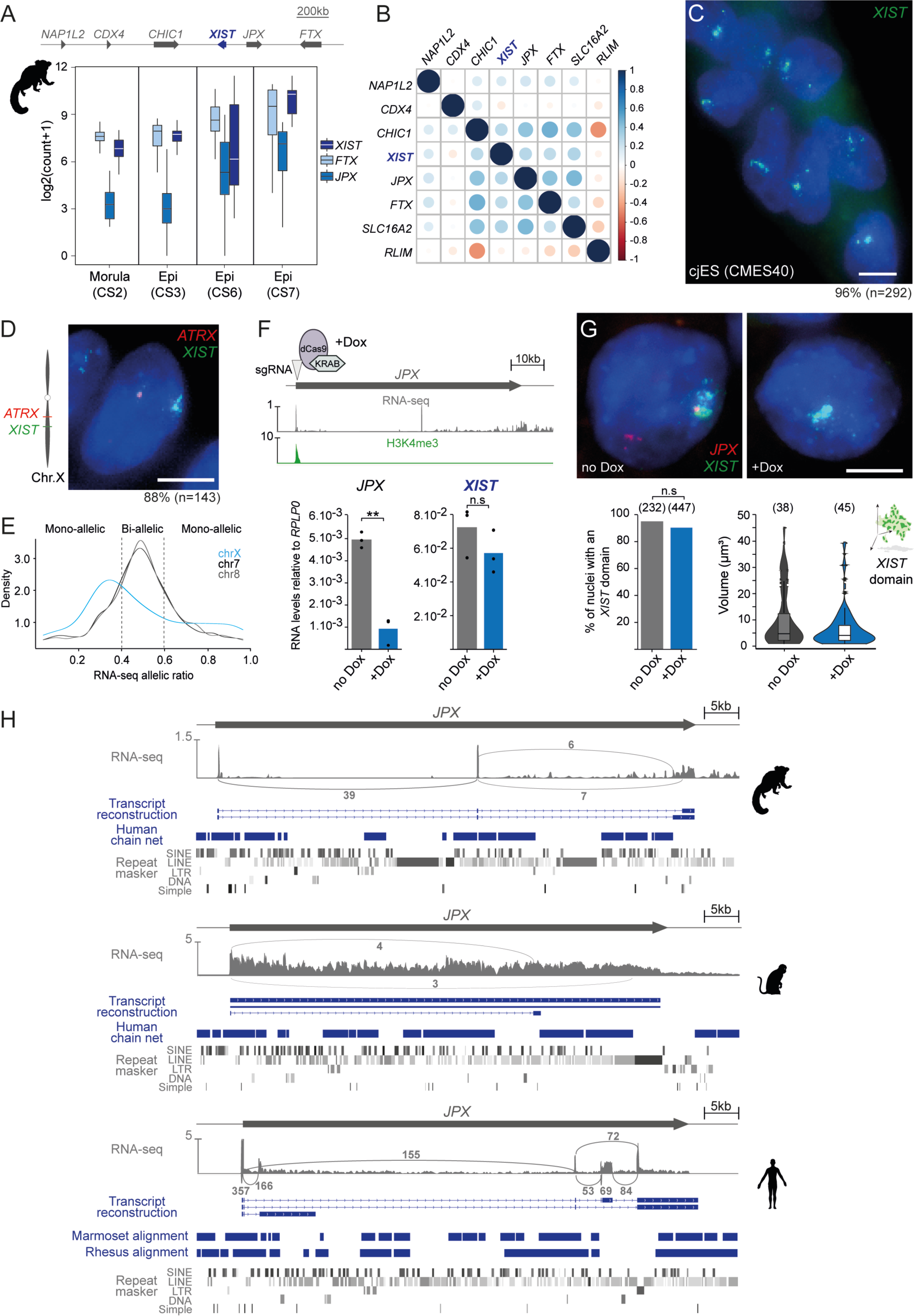
*JPX* inhibition does not affect *XIST* expression in cjESCs. A. *XIST*, *JPX* and *FTX* expression dynamics during pre-implantation development of female white-tufted-ear marmoset extracted from RNA-seq data from^92^. B. Pearson correlation between *XIC* gene expression profiles during pre-implantation development of female white-tufted-ear marmoset computed using RNA-seq data from^92^. C. Representative image of *XIST* RNA-FISH in CMES40 cjESCs. The percentage of nuclei with an *XIST* RNA accumulation is indicated. (Scale bar: 10 µm). D. Representative image of double RNA-FISH for *XIST* and for *ATRX* in CMES40 cjESCs. The percentage of cells showing monoallelic *ATRX* transcription is indicated. (Scale bar: 5 µm). E. Density plots of allelic ratios at polymorphic positions on chromosome X, 7 or 8 extracted from CMES40 cjESCs RNA-Seq data (SRR11996369). F. Dox-inducible CRISPRi strategy showing the position of the guide RNA targeting the *JPX* promoter, RNA-Seq profile and H3K4me3 CUT&RUN distribution along the *JPX* transcription unit of CMES40 cjESCs. Below, average expression levels of *JPX* and *XIST* measured by RT-qPCR in *JPX* CRISPRi cjESCs treated (+dox) or not (no dox) with doxycycline (n=3). Unpaired two-sided t-test: n.s., not significant; **p-value < 0.01. G. Representative images of *XIST* RNA-FISH in *JPX* CRISPRi cjESCs treated (+dox) or not (no dox) with doxycycline. The percentages of cells with an *XIST* cloud are shown on the bar-plot aside. Fisher test: n.s., not significant. Violin plots show the distributions of the volumes of *XIST* RNA accumulations. Wilcoxon test: n.s., not significant. Scale bar = 5 µm. H. Transcription profiles of marmoset, macaque and human *JPX* orthologs in CMES-40 cjESCs, LYON-ES1 rhESCs and H9 hESCs respectively. Arcs represent the number of reads supporting the splicing junctions reconstructed from RNA-Seq data using the Scallop program (see Methods). Sequence alignments from the UCSC browser show conservation blocks in blue as well as the position of repeated sequences (RepeatMasker annotation) along each ortholog.

We used a *Callithrix jacchus* ESC (cjESC) line CMES40^65^, the transcriptome of which resembles that of marmoset post-implantation epiblast (Supplementary Fig. 6A). CjESCs strongly and homogeneously express the pluripotency marker *POU5F1* (Supplementary Fig. 6B), display a *XIST* domain in 96% of cells (Fig. 4C) and focal accumulations of H2AK119ub in 63% of nuclei (Supplementary Fig. 6C). These accumulations colocalise with H3K27me3 enrichment in only 13% of cells (Supplementary Fig. 6C), suggesting specific dynamics of H3K27me3 deposition in this species that need to be confirmed *in vivo*. Mono-allelic expression of the X-linked gene *ATRX* away from the *XIST*-coated X (Fig. 4D) and an X-linked allelic ratio of 0.3 (Fig. 4E) confirmed that CMES40 cjESCs, like primed rhESCs and hESCs, have undergone XCI. This cjESC line thus model’s similar developmental stages as the LYON-ES1 rhESC and H9 hESC used in this study.

In cjESCs, like in rhESCs, *JPX* escapes from XCI in 59% of cells, as measured by RNA-FISH (Supplementary Fig. 6D). To interrogate the function of *JPX* in *XIST* regulation in cjESC, we knocked-down *JPX* in an inducible manner, as previously described for rhESCs (Fig. 4F). Transgene induction resulted in efficient repression of *JPX* transcription, as measured by RT-qPCR (Fig. 4F) and by RNA-FISH (Supplementary Fig. 6E). No significant differences in *XIST* RNA levels (Fig. 4F), in the percentage of cells showing an *XIST* accumulation (Fig. 4G) or in the volume of *XIST* domains (Fig. 4G) between induced cells and non-induced cells could be detected. This indicates that *JPX* is not a significant component of the *XIST* regulatory network in cjESCs.

Examination of the transcript structures of the *JPX* orthologs expressed in female cjESCs, rhESCs and hESCs revealed wide divergences in the DNA sequence (shown by the discontinuous Chain net alignments), exonic structures and accumulation of TE insertions (RepeatMasker) (Fig. 4H). More precisely, there is no conservation between species for splicing junctions and no apparent selective constraint for a spliced *JPX* molecule in primates, as highlighted by the lack of splicing of macaque *JPX* and the three-exon structure that results from the insertion of a SINE element in the middle of the marmoset transcription unit (Fig. 4H). This contrasts with the *XIST* orthologs for which DNA sequence and transcript structure are mostly conserved between the three primate species we studied (Supplementary Fig. 6F). Such a lack of conservation is indicative of the absence of selective pressure on *JPX* RNA products and suggests that these molecules do not mediate *JPX* function in *XIST* regulation in rhESCs and cjESCs. In addition, transcription levels across the *JPX* unit in cjESCs are markedly low compared to levels in rhESCs or to hESCs (Fig. 4H), further suggesting that *JPX* transcription does not significantly contribute to *XIST* regulation in marmosets.

These analyses support a gradual recruitment of *JPX* to *XIST* regulation during primate diversification: in human, *JPX* transcriptional activity controls *XIST* expression profiles as previously described^41^; in rhesus, a similar mode of action is likely to operate but *JPX* contribution to *XIST* expression is reduced; in marmoset, neither *JPX* RNAs nor *JPX* transcription seem to play a significantly role in *XIST* regulation.

### Enh5 and *JPX* exert additive effects on *XIST* regulation in rhesus ESCs

CUT&RUN analyses of *XIC* chromatin states in rhESCs and hESCs revealed the presence of two candidate cRE, Enh3 and Enh5, that are enriched in H3K27ac and H3K4me1 and fall into open chromatin regions in macaque but not in human cells (Fig. 5A). Hence, these entities may exert enhancer functions in rhESCs specifically. Since many enhancers physically engage with target promoters^6–8^, we used the capture HiC data to probe for interactions between Enh3/5 and *XIST* promoter in LYON-ES1 rhES and H9 hESCs. Reciprocal 4C analyses detected significant contact frequencies between *XIST* transcriptional start site (TSS) and Enh5 but not with Enh3 in rhESCs (Supplementary Fig. 7A). Conversely, in hESCs, no interaction with either Enh3 or Enh5, classified as poised enhancers in these cells, could be detected (Supplementary Fig. 7B).

**Figure 5:**
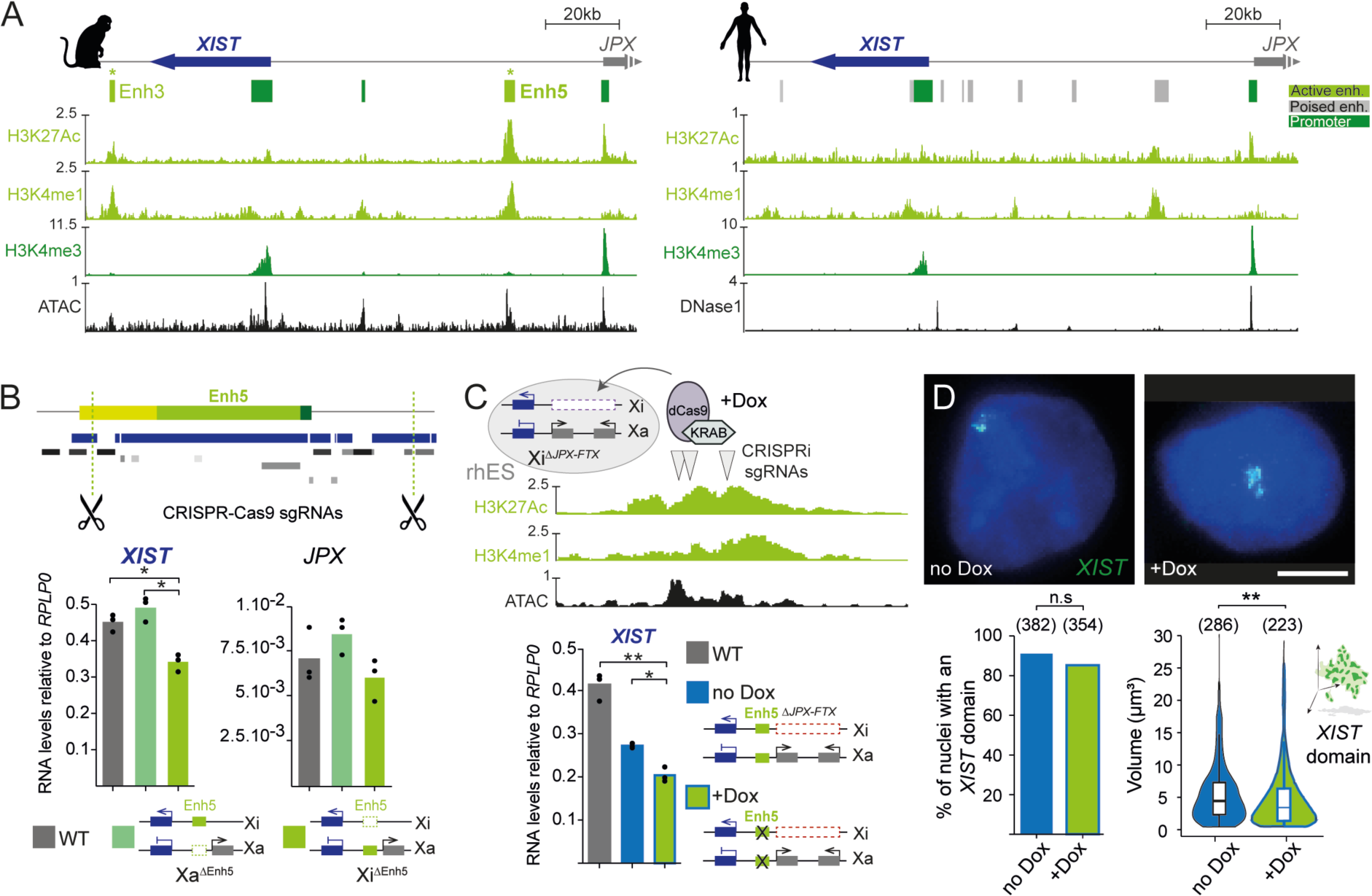
Enhanced effect of Enh5 decommission on *XIST* levels in τ<*JPX-FTX* rhESCs. A. CUT&RUN distributions of H3K27ac, H3K4me1, H3K4me3 and ATAC-seq profiles along the indicated region in LYON-ES1 rhESC (this study, left) and ChIP-seq distributions of the same histone modifications and DNase I sensitive sites in H9 hESCs (ENCFF291ZMA, right). B. Enh5 CRISPR-Cas9 deletion strategy in rhESCs showing the position of the guide RNAs with respect to repeats (repeatMasker annotation, grey shades) and to blocks of sequence conservation with human (Human chain net, dark blue). Below, average expression levels of *XIST* and *JPX* measured by RT-qPCR in WT and in clones in which the enhancer 5 has been deleted on the Xa (Xa^ΔEnh5^) or on the Xi (Xi^ΔEnh5^) (n=3). Dots represent expression levels in independent clones. Unpaired two-sided t-test: *p-value < 0.05. C. Inducible CRISPRi strategy used to decommission the Enh5 in the context of the *JPX-FTX* deletion (Xi^ΔJPX-FTX^ rhESCs). The positions of the guide RNAs (arrowheads) relative to H3K27ac, H3K4me1 and ATAC-Seq peaks is shown. *XIST* expression levels measured by RT-qPCR in WT and in Enh5 CRISPRi Xi^ΔJPX-FTX^ rhESCs treated (+Dox) or not (no Dox) with doxycycline. Unpaired two-sided t-test: *p-value < 0.05; **p-value < 0.01. D. Representative images of *XIST* RNA-FISH in rhESCs deleted for *JPX-FTX* on the Xi (no Dox) and upon Enh5 decommission (+Dox). The percentages of cells with a *XIST* cloud are shown on the bar-plot underneath. Fisher test: n.s., not significant. Violin plots show the distributions of the volumes of *XIST* RNA accumulations. Wilcoxon test: n.s., not significant; **p-value < 0.01. Scale bar = 5 µm.

Deleting Enh3 using CRISPR-Cas9 technology in LYON-ES1 rhESCs did not change *XIST* expression levels (Supplementary Fig. 7C) or the expression of any other genes of the region (Data not shown). We then repeated this approach to generate heterozygous deletions of Enh5 (Fig. 5B and Supplementary Fig. 7D). *XIST* RNA levels as measured by RT-qPCR appeared significantly reduced (∼1.4 fold) when the deletion was located on the Xi compared to WT cells or to Xa^ΔEnh5^ clones while other *XIC* genes like *JPX* remained unaffected (Fig. 5B). This indicates that Enh5 specifically contributes to *XIST* expression on the Xi.

The impact of Enh5 deletion on *XIST* expression is however quite mild, which suggests that different molecular actors, including *JPX,* may exert additive or synergistic effects on *XIST* regulation as described at other loci^66–68^. To test this hypothesis, we engineered an inducible decommission of Enh5 in LYON-ES1 rhESCs carrying the *JPX-FTX* deletion on the Xi using Dox inducible CRISPRi (Fig. 5C). As expected, CRISPRi induction triggered the targeted deposition of H3K9me3 and depletion of H3K27Ac over the Enh5 (Supplementary Fig. 7E). Under this condition, *XIST* transcript levels appeared reduced by ∼1.4 fold compared to cells in which Enh5 is still active and by ∼2 fold compared to WT cells (Fig. 5C). RNA-FISH analysis, however did not show any significant reduction in the percentage of nuclear *XIST* accumulations upon Enh5 decommission in Xi^Δ*JPX*-*FTX*^ cells (+Dox), only a slight decrease in the volume of *XIST* nuclear domains was detected (Fig. 5D).

Altogether this indicates that *JPX* and Enh5 exert additive regulatory effects on *XIST* expression levels but do not on control the probability of *XIST* to coat the Xi of rhESCs.

### *XIST* transcriptional regulation follows the Constructive Neutral Evolution hypothesis

To determine whether functionality and sequence conservation are connected in the case of *XIST* non-coding regulators in primates, we analysed the mode of selection of the CTCF sites within the macaque HERVK, the Enh5 cRE defined in macaque and the *JPX* transcription start site (TSS). A measure of the evolution rate of DNA sequences is the phyloP score. Any given stretch of nucleotides that encodes a function with a positive effect on the organism’s fitness should have, when mutated, a deleterious effect for the host. Hence, nucleotides of such a sequence should be under constraint, ie. more conserved between species than expected under neutral drift (positive phyloP score). Purifying or directional selection are characterised by slower or accelerated substitution rates, respectively, compared to neutrally evolving sites. The statistical test package in the phyloP program computes divergence from neutrality at the nucleotide level and outputs positive values for constrained sites and negative values for sites under accelerate substitution rates^69^.

Here we worked with the most recent phyloP scores calculated from multiple alignments of 241 mammalian species including 42 primates and 16 cercopithecidae^2^. We used either the human (Hg38) or the rhesus macaque (rheMac10) as reference genomes. According to this analysis, the HERVK CTCF motifs have neutral phyloP scores (Fig. 6A). The Enh5 cRE is also evolving neutrally in both humans and macaques (Fig. 6B). In contrast, the *JPX* TSS show distinct evolutionary behaviours in humans and macaques: the human TSS displays a neutral phyloP score while, in macaques, the negative phyloP score marks accelerated substitution rates and indicates directional selection potentially leading to the emergence of a new function (Fig. 6C). This is in sharp contrast with the *XIST* TSS, which is under strong selective constraint in both humans and macaques, consistent with its essential role during mammalian development.

**Figure 6:**
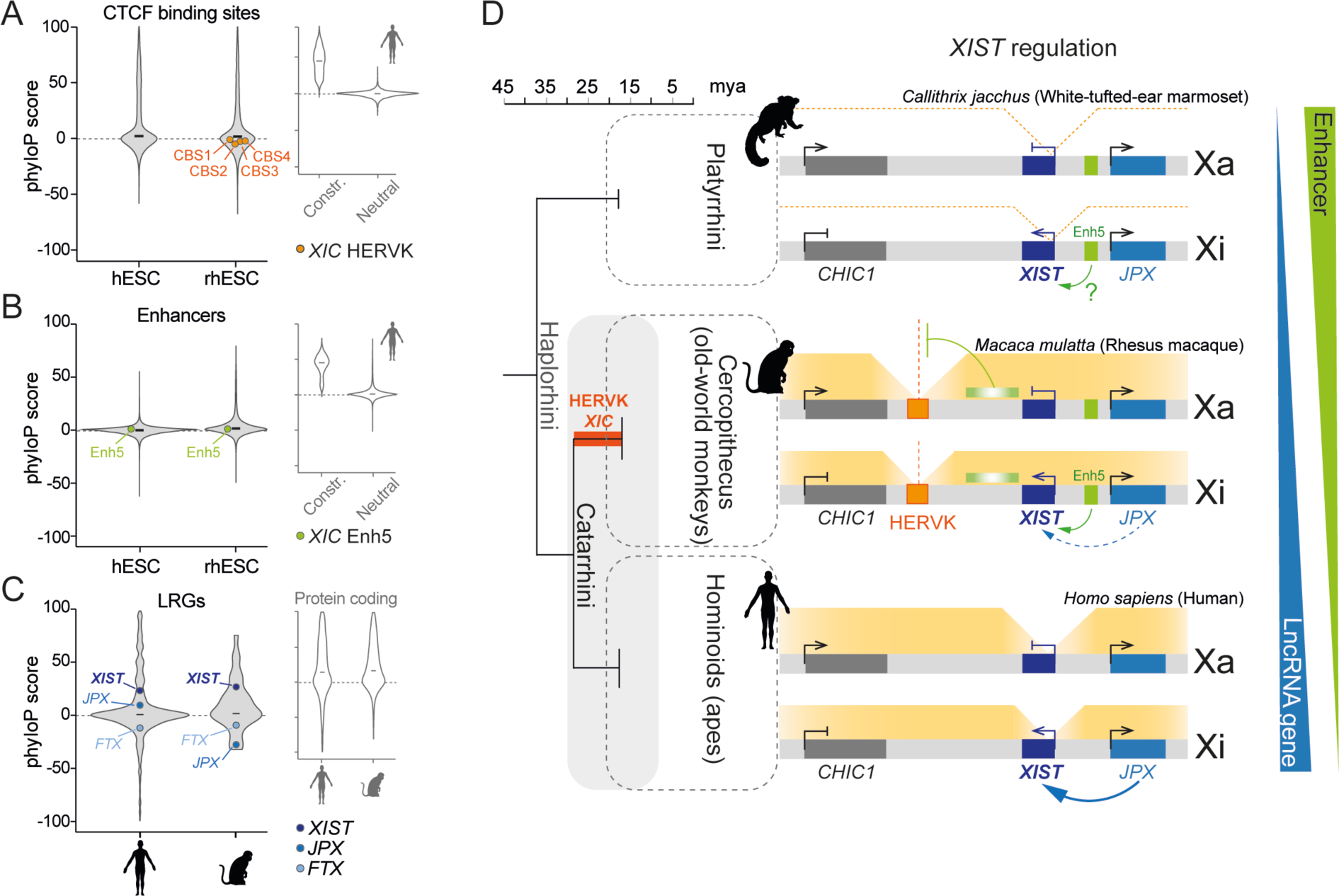
Evolution of *XIST* regulatory modalities in primates. A. Distribution of phyloP scores for CTCF binding sites linked to chromosomes 7, 8 and X in H9 hESCs extracted from ChIP-seq data (ENCODE) (n= 3160) or in LYON-ES1 rhESCs extracted from CUT&RUN analyses (n= 6865). The position of HERVK embedded CTCF binding sites within the rhESC distribution is indicated. As a comparison the distribution of constrained and not constrained CTCF motifs as annotated by the Zoonomia consortium^16^ in H9 hESCs is shown on the top right violin plots. B. Same as panel A for sequences with active enhancer signatures. Enh5 follows a neutral evolution regimen. C. Same as panel A for LRG transcription start sites (TSS). TSS annotations from the FANTOM5 consortium have been used for both species. TSS of protein coding genes are shown for comparison. D. Proposed evolution of the *XIST* regulatory network in primates inferred from the present study and from previous observations^41^.

None of the regulatory elements we studied here are under selective constraint. We have however carefully demonstrated their involvement in *XIST* regulation. These observations support the Constructive Neutral Evolution (CNE) theory^22,26^. According to CNE, some sequences promote cellular and molecular complexity without leading to a fitness boost or loss for the population and are thus evolving neutrally. The functional innovations they provide can however become selected in the population upon environmental changes. Our results suggest that the non-coding players of the *XIST* regulatory network in primates evolve under CNE, thereby providing some of the first experimental data that support this theory in the frame of specific biological processes.

## Discussion

Characterising the evolutionary modalities of biological processes and identifying the genomic elements more prone to provide species with adaptative potentials represent important challenges of modern biology. So far, studies that attempted to connect changes in a given regulatory landscape to functional outcomes are scarce and only consider large evolutionary timescales that allow capturing drastic regulatory variations but not subtle or transitory innovations. In addition, many non-coding regulators are poorly conserved, which calls for dedicated approaches that do not rely on sequence comparison. To this end we generated high-resolution capture-HiC maps of the *XIC* in ESCs from closely-related rhesus macaques and humans that we crossed with the distributions of chromatin features typical of regulatory elements. This revealed that, over only 35 mya of evolution, *XIST* regulatory landscapes have significantly diverged in both 3D topology and molecular mechanisms. Capture-HiC maps generated from chimp iPSCs (chimps diverged from humans 5 mya) appeared superimposable with hESC’s maps (Supplementary Fig. 8), indicating that species evolutionary closer to humans do not provide additional information. These results validate screens for regulatory innovations based on interspecies comparisons of functional maps as powerful approaches to efficiently trace the evolutionary trajectory of a given gene regulatory system.

One striking finding is the emergence of a HERVK-driven chromatin boundary that expands *XIST* interaction domain centromerically in rhESCs. TEs actively contribute to the short-term renewal of transcription factor binding sites^16^; HERVK and HERVH in particular are strongly associated with species-to-species divergences in chromatin architecture^70–72^. However, the regulatory consequences of such evolution are globally unknown. Deleting the *XIST* HERVK reproduces the topological configuration of the human *XIST* locus but does not significantly impact *XIST* expression on the Xi, at least in the context of ESCs. Instead, it induces a modest up-regulation of centromeric genes *NAP1L2* and *CHIC1* on the Xa that have become potential targets of Enh3. This rather mild effect may have different interpretations. First, the HERVK insertion may have a neutral impact on the fitness of the species.

Yet, the element is present in various species of Cercopithecidae and most CTCF binding sites that confer the barrier activity are conserved in these species. Alternatively, the reshaping of the chromatin topology in response to changes in the regulatory network may be required to circumvent the establishment of enhancer-promoter pairs that may be deleterious in other cellular contexts. Such cell type specific effects have been recently illustrated by the insertion of the *Zfp42* gene into the highly structured regulatory environment of the *Fat1* gene in the mouse^73^. We did not observe any changes in cell morphology, metabolism or proliferation upon HERVK mutation in rhESCs. However, other cell identities or cell type transitions need to be tested.

We also showed that the function of *JPX* as an *XIST* regulator progressively diversified in primates: from no detectable regulatory activity in the marmoset (Platyrrhini) to a mild effect that only impacts *XIST* RNA levels in macaques (Catarrhini; Cercopithecus) and a stronger contribution that impacts both *XIST* RNA levels and the percentage of cells with an *XIST* domain in humans (Catarrhini; Hominoids)^41^ (Fig. 6D). In parallel, the Enh5 cRE is recruited to *XIST* regulation in rhESCs and probably also in marmosets since the orthologous sequence is enriched in H3K27Ac and depleted of H3K9me3 in cjESCs (Supplementary Fig. 7F). These observations suggest two putative scenarios for the evolution of *XIST* regulatory network that are equally parsimonious. In the first scenario, *XIST* is regulated by Enh5 in the primate common ancestor and *JPX* is inoperative (case of the marmoset), *JPX* function is then acquired in Catarrhini and reinforced in Hominoids in which Enh5 becomes inactive. This scenario implies a convergent evolution of *JPX* as a *XIST* regulator in mice and primates that is mediated by different modules of the *JPX* LRG: *Jpx* RNA in mice and *JPX* transcription in humans^39,41,62^. In the second scenario, *JPX* initially regulates *XIST* in the mice and primate common ancestor but functional implication is lost independently in marmosets and macaques. Studies of *JPX* function in Dermopteras (flying lemurs), the closest primate relatives, would allow to define the regulatory modalities in the rodent-primate common ancestor and distinguish between these two hypotheses.

XCI is essential to female embryo development and therefore represents a biological process under strong purifying selection. *XIST* lncRNA, the key player of the process in placental mammals, is well conserved in sequence and structure, which indicates that it is subject to significant selective pressure. In contrast, we did not find evidence of ongoing transcription at syntenic positions of *Tsix*, *Linx* and *Xite* LRGs that are prominent *Xist* repressors in mice. In addition, *JPX*, a positive regulator of *XIST* in mice and humans, harbours extensively reshuffled transcript structures in primates and a promoter sequence that is not conserved or experience accelerated substitution rates in humans and macaques respectively. Likewise, Enh5 and rhesus HERVK both lack sequence conservation. In primate ESCs, these LRG and cREs have incremental effects on *XIST* regulation and are unlikely to significantly impact the maintenance of XCI since the silencing of X-linked *ATRX* and *POLA1* is maintained upon mutation of these elements. Altogether, this suggests that these non-coding actors follow the Constructive Neutral Evolution (CNE) regimen^22,26^, which predicts that novel functionalities evolve in a host with no consequence on the fitness of the population but that may lead to a fitness gain later on, upon changes in selective pressure parameters. Therefore, non-coding players of the *XIC* may provide the XCI process with a reservoir of adaptative potentials. Importantly, our functional studies are among the first to support CNE in the context of a specific biological process. Additional analyses at the level of primate populations would be required to validate the CNE hypothesis in the frame of XCI evolution.

Are there other *XIST* regulators either within the *XIC* or outside, like *trans*-acting transcription factors, in primates? Conserved contribution of transcription factors such as YY1 might ensure robustness to *XIST* expression since regulatory sequences tend to be encoded within the promoter region of essential genes^34^. In this case, cREs and LRGs would serve to integrate environmental cues and slightly modulate *XIST* expression dynamics. These actors seem to regulate *XIST* differently in primate species. Indeed, we noticed that human *JPX* controls both *XIST* probability to be expressed and *XIST* RNA levels, while the combined action of rhesus *JPX* and Enh5 only impacts *XIST* expression levels. It is interesting to note that this shift of mechanisms correlates with a change in *XIST* expression stability during prolonged cell culture: hESCs tend to lose *XIST* expression over passages as part of a process called XCI “erosion”^74,75^ while *XIST* RNA accumulation on the Xi appears highly stable in rhESCs or in cjESCs (data not shown). Hence the effects of *XIST* regulatory modalities may only be revealed under specific circumstances. More generally, remodelling of *XIST* regulators may reflect cell type or developmental stage specificities/requirements in each primate species. Indeed, XCI kinetics in macaque and human embryonic and extra-embryonic tissues seem to differ significantly^46^. In conclusion, this study traces the evolutionary trajectory of *XIST* regulators in primates and reveals how non-adaptive evolution likely explains the plasticity of gene regulatory networks in mammals.

## Supplementary figure legends

**Supplementary Figure 1: Rhesus macaque iPSCs are in a primed state of pluripotency with established X-inactivation**

A. Heatmap of Pearson correlation between LYON-ES1 rhESCs transcriptome^54^ (two independent replicates) and single cell RNA-seq data from cynomolgus macaque early development^46,52^. Correlation is computed on the expression of the top 40 genes defining each epiblast stage.

B. *XIST* RNA-FISH in female rhesus macaque iPSC^55^. The percentage of nuclei with a *XIST* RNA accumulation is indicated. (Scale bar: 5 µm).

C. H2AK119Ub or H3K27me3 immunofluorescence staining of rhesus macaque iPSCs. The percentages of nuclei with H2AK119Ub or H3K27me3 focal enrichments are indicated. (Scale bar: 10 µm)

D. Triple RNA-FISH for *XIST* (green), *ATRX* (white) and *POLA1* (red) in rhesus macaque iPSCs. Gene positions on the macaque X chromosome are indicated. The bar plot shows the percentages of indicated expression patterns in the cell population. (Scale bar: 10 µm).

**Supplementary Figure 2: *XIC* chromatin landscapes in rhECS or hESCs**

A. Coverage of capture probes targeting human (top) and macaque (bottom) *XIC*s.

B. Pearson correlation between capture HiC replicates performed on hESCs (right) and or on rhESCs (left) with 6 kb bins.

C. CUT&RUN distribution of H3K27Ac, H3K4me1 and H3K4me3 and of ATAQ-seq accessible chromatin regions along the *XIC* in LYON-ES1 rhESCs. Histone mark peaks as determined by MACS2 (see Methods) are displayed below.

D. Same as panel C for H9 hESCs. DNAse I-seq ENCODE data (ENCFF291ZM1) were used to map chromatin accessibility.

**Supplementary Figure 3: HERVK sequence conservation across cercopithecidae**

A. Multiple alignments of the *XIC* HERVK sequence from representative species of the Cercopithecus family, represented in the phylogenetic tree, on the *Macaca mulatta (mm)* HERVK sequence. Alignment color code is indicated below.

B. Close-up on CTCF binding site region. Arrows depict the position and direction of CTCF binding sites (CBS) numbered from one to four, from the 5’ to 3’ of the DNA sequence. Nucleotide substitutions are shown.

**Supplementary Figure 4: Characterization of the CRISPR-Cas9-mediated HERVK deletion in LYON-ES1 rhESCs**

A. Sequencing results of PCR products (primer positions are shown in orange) encompassing the HERVK deletion from three independent LYON-ES1 rhESC clones carrying the deletion on the Xa (Xa^ΔHERVK^) and from three rhESC clones carrying the deletion on the Xi (Xi^ΔHERVK^) that have been used in this study. The dotted orange line indicates the theoretical position of the junction upon deletion. The star indicates the polymorphism used for allelic assessment of the deletion. Primers used to amplify the polymorphic region prior to sequencing are shown in grey.

B. Allelic assessment of HERVK deletion in rhESC clones shown in A. Chromograms show the sequence of intact alleles at the polymorphic position indicated in panel A in independent WT rhESC clones and in deleted clones.

C. Average expression levels of pluripotency markers *NANOG* and *POU5F1* measured by RT-qPCR in WT, Xa^ΔHERVK^ or Xi^ΔHERVK^ rhESC clones (n=3). Each dot represents the expression levels in independent clones. Unpaired t-test did not detect any significant difference between the genotypes.

D. Average expression levels of *SCL16A2* and *RLIM* measured by RT-qPCR in WT, Xa^ΔHERVK^ or Xi^ΔHERVK^ rhESC clones (n=3). Dots represent the expression levels in each clone. Unpaired t-test did not detect any significant difference between the genotypes.

E. Virtual 4C profiles showing the distribution of interaction frequencies along the depicted region taking *CHIC1* TSS (top) or *NAP1L2* TSS (bottom) as viewpoints in WT LYON-ES1 rhESCs (grey) and in rhESCs carrying the HERVK homozygote deletion (orange).

**Supplementary Figure 5: Characterization of CRISPR-Cas9 deletion of the *JPX-FTX* locus in rhESCs**

A. Sequencing results of PCR products (primer positions are shown in blue) encompassing the *JPX-FTX* deletion from three independent LYON-ES1 rhESC clones carrying the deletion on the Xa (Xa^ΔJ*PX-FTX*^) and from the three rhESC clones carrying the deletion on the Xi (Xi^ΔJ*PX-FTX*^) that have been used in this study. The blue dotted line indicates the theoretical position of the junction upon deletion. The star indicates the polymorphism used for allelic assessment of the deletion. The primers used to amplify the polymorphic region prior to sequencing are shown in grey.

B. Allelic assessment of *JPX-FTX* deletion in rhESC clones depicted in A. The chromograms show the sequence of intact alleles at the polymorphic position indicated in panel A in independent WT rhESC clones and in deleted clones.

C. Average expression levels of pluripotency markers *NANOG* and *POU5F1* measured by RT-qPCR in WT, Xa^ΔJ*PX-FTX*^ and Xi^ΔJ*PX-FTX*^ rhESC clones (n=3). Each dot represents the expression levels in independent clones. Unpaired t-test did not detect any significant difference between the genotypes.

D. Representative image of triple RNA-FISH for *XIST* (green), *JPX* (red) and *FTX* (white) in LYON-ES1 rhESCs carrying the *JPX-FTX* deletion on the Xa (top) or on the Xi (bottom). Percentages of indicated transcription patterns are shown on the bar plot. (Scale bar: 5 µm).

E. Violin plot showing the distribution of volumes of *XIST* RNA accumulations in WT rhESCs or in Xi^ΔJ*PX-FTX*^ cells. Wilcoxon test: **p-value < 0.01.

F. Triple RNA-FISH for *XIST* (green), *ATRX* (white) and *POLA1* (red) in WT LYON-ES1 rhESCs and in Xi^ΔJ*PX-FTX*^ cells. The position of the genes on the macaque X chromosome is shown. The percentages of cells with depicted transcription patterns are shown on the bar-plot. (Scale bar: 5 µm).

G. Table showing the number of insertions of the rTA/sgRNA expressing transgene and of dCas9-KRAB transgene in each LYON-ES1 clone determined by qPCR.

H. Double RNA-FISH for *XIST* and for *JPX* upon doxycycline induction of the CRISPRi system targeting the *JPX* promoter in LYON-ES1 rhESCs. The percentages of cells with a *XIST* cloud are shown on the bar-plot aside. Chi-square test: ****p-value < 0.0001.

**Supplementary Figure 6: Characterization of CMES40 cjESCs**

A. Heatmap of Pearson correlation between CMES40 cjESCs transcriptome (GSM4610404) and single cell RNA-seq data from the epiblast or from embryonic disk stages of marmoset development^92^. Correlation is computed on the expression of the top 40 genes defining each stage.

B. Representative immunofluorescence staining of pluripotency factors POU5F1 in CMES40 cjESCs. (Scale bar: 10 µm).

C. Representative H2AK119Ub and H3K27me3 co-immunofluorescence staining in CMES40 cjESCs. The percentages of cells with foci are indicated below. (Scale bar: 5 µm).

D. Representative image of double RNA-FISH for *XIST* (green) and for *JPX* (red) in CMES40 cjESCs. The percentage of cells with bi-allelic transcription of *JPX* is indicated below. (Scale bar: 5 µm).

E. Representative pictures of double RNA-FISH for *XIST* (green) and for *JPX* (red) in *JPX* CRISPRi cjESCs treated (+Dox) or not (no Dox) with doxycycline. Percentages of cells with the indicated transcription pattern are shown on the bar plots. Chi-square test: ****p-value < 0.0001.

F. Transcription profiles of marmoset, macaque and human *XIST* orthologs in CMES40 cjESCs, LYON-ES1 rhESCs and H9 hESCs, respectively. Sequence alignments from the UCSC browser show conservation blocks in grey as well as the position of repeated sequences (RepeatMasker annotation) along each ortholog.

**Supplementary Figure 7: Characterization of Enh3 and Enh5 deletions in rhESCs**

A. Virtual 4C profiles showing the distribution of interaction frequencies along the depicted region taking Enh3 (top), Enh5 (middle) or *XIST* TSS (bottom) as viewpoints in LYON-ES1 rhESCs. The arcs underneath show enhancer-promoter predicted interactions (ABC model)^8^.

B. Same as panel A in H9 hESCs.

C. CRISPR-Cas9 deletion of Enh3 cRE and *XIST* expression levels as measured by RT-qPCR in WT LYON-ES1 rhESCs and in cells carrying a homozygous deletion of Enh3.

D. Schematic representation of the CRISPR-Cas9 knock out of the Enh5 candidate cRE in LYON-ES1 rhESCs. The chromograms show the sequence of intact alleles at the polymorphic position indicated in independent WT rhESC clones and in deleted clones.

E. ChIP for H3K9me3 and for H3K27Ac in LYON-ES1 upon Enh5 decommission (+Dox) compared to non-decommissioned cells (no Dox) followed by qPCRs quantification at indicated positions along the Enh5 locus.

F. Same as panel E along Enh5 orthologous region in CMES40 cjESCs.

**Supplementary Figure 8: Capture HiC map of the XIC of Chimp iPSCs**

A. Capture-HiC maps of the *XIC* span in chimp-iPSCs showing interaction frequencies calculated over 6-kb bins (n=2). The distribution of insulation scores and coverage of capture probes are shown underneath.

B. Pearson correlation between capture HiC replicates performed on chiPSC Sandra^93^, with 6 kb bins.

## Summary of Supplementary Tables

**Supplementary Table 1:** List of DNA oligos targeting rhesus macaque sequences used in this study.

**Supplementary Table 2:** List of DNA oligos targeting marmoset sequences used in this study.

**Supplementary Table 3:** List of antibodies used in this study.

**Supplementary Table 4:** Accession number for publicly available datasets used in this study.

**Supplementary Table 5:** List of BAC DNA used for RNA-FISH probes generation in this study.

## Methods

### Primate pluripotent stem cell lines and culture conditions

Female LYON-ES1 rhESCs^51^ were grown on gelatin-coated dishes with feeder cells (mouse embryonic fibroblasts; 18000/cm^2^) in KO-DMEM (Gibco-Thermo Fisher Scientific) containing 20% of KO serum (Gibco-Thermo Fisher Scientific) complemented with Penicillin-Streptomycin-Glutamine (1XPSG; Gibco-Thermo Fisher Scientific), with non-essential amino acids (1XMEM Non-Essential Amino Acids Solution; Gibco-Thermo Fisher Scientific), with 100 μM of beta-Mercaptoethanol (Gibco-Thermo Fisher Scientific) and with 5 ng/mL of basic-FGF (Gibco-Thermo Fisher Scientific). Cells were routinely passaged as clumps using 1mg/ml Collagenase IV (Gibco-Thermo Fisher Scientific). For experiments requiring single-cell suspension, cells were incubated with Accutase (Stemcell Technologies) and plated in fresh rhESC media supplemented with 10 μM of Y-27632 (Stemcell Technologies).

The rhesus macaque iPSC line rhiPSC24, was cultured as previously described in^55^. Briefly, rhiPSC were grown on Geltrex-coated culture dishes (Gibco-Thermo Fisher Scientific) in StemMACS iPSC-Brew XF (Miltenyi Biotec) supplemented with 0.5 µM CHIR99021 (Axon medChem) and 1 µM IWR1-endo (Merck). RhiPSC24 were routinely passaged in clumps using Gentle Cell Dissociation reagent (Stemcell Technologies) according to the manufacturer instructions.

Female CMES40 cjESCs^65^ were grown on gelatin-coated dishes with feeder cells (mouse embryonic fibroblasts; 14000/cm^2^) in KO-DMEM (Gibco-Thermo Fisher Scientific) containing 20% of KO serum (Gibco-Thermo Fisher Scientific) complemented with Penicillin-Streptomycin-Glutamine (1XPSG; Gibco-Thermo Fisher Scientific), with non-essential amino acids (1XMEM Non-Essential Amino Acids Solution; Gibco-Thermo Fisher Scientific), with 100 μM of beta-Mercaptoethanol (Gibco-Thermo Fisher Scientific) and with 8 ng/mL of basic-FGF (Gibco-Thermo Fisher Scientific). Cells were routinely passaged as single cell using TrypLE (Gibco-Thermo Fisher Scientific).

Female H9 hESCs were obtained from the WiCell Research Institute and cultured on Matrigel-coated culture dishes in mTeSR™1 media (Stemcell Technologies) according to the manufacturer instructions. They were routinely passaged in clumps using gentle cell dissociation reagent (Stemcell Technologies) according to the manufacturer instructions. For experiments requiring single-cell suspension, cells were incubated with Accutase (Stemcell Technologies) and plated in fresh mTeSR™1 media supplemented with 10 μM of Y-27632 (Stemcell Technologies).

Female chiPSCs^76^ were grown as H9 hESCs.

All cell lines were grown in 5% O2 and 5% CO2 at 37°C.

Research on human embryonic stem cells has been approved by the Agence de la Biomédecine and informed consent was obtained from all subjects.

### RNA FISH and Immunofluorescence staining

For RNA-FISH, cells grown on coverslips overnight were fixed in 3% paraformaldehyde (Electron Microscopy Science) for 10 min, permeabilized in CSK buffer (10 mM PIPES; 300 mM sucrose; 100 mM NaCl; 3 mM MgCl2; pH 6.8) supplemented with 0.5% Triton X-100 (Sigma-Aldrich) and 2 mM VRC (New England Biolabs) for 7 min on ice and stored in 70% ethanol. For probe preparation, 1 μg of fosmid or BAC DNA was labelled by nick-translation with fluorescent dUTPs (Abott Molecular or GE HealthCare Life Science) for 3 h at 15°C as described previously^41^. Hybridization mix containing 100 ng of each probe supplemented with 1 μg of Cot-I DNA (Invitrogen) and 3 μg of Sheared Salmon Sperm DNA (Invitrogen) in deionized formamide (Sigma Aldrich) was denatured for 10 min at 75°C, mixed with an equal volume of 2XHybridization Buffer (4XSSC, 20% dextran sulfate, 2 mg/ml BSA, 2 mM VRC) and hybridized on ethanol dehydrated cells overnight at 37°C in a humid chamber. Coverslips were then washed three times in 50% formaldehyde/2XSSC (pH 7.2) for 4 min at 42°C, three times in 2XSSC for 4 min at 42°C and mounted in Vectashield plus DAPI (Vector Laboratories).

For immunofluorescence staining, cells were prepared and fixed as described above then stored in PBS1X. Just before staining, cells were permeabilised in ice-cold CSK 0.5% Triton X-100 for 7 min. Cells were blocked in 1XPBS/1% BSA (Sigma-Aldrich) for 15 min at RT, incubated for 45 min at RT with primary antibody diluted in 1X PBS/1% BSA (H3K27me3, Active motif-61017; H2AK119Ub, Cell signalling-8240S), washed with 1XPBS then incubated for 40 min at RT with Alexa Fluor 488 nm anti-rabbit or Alexa Fluor 594 nm anti-mouse secondary antibodies (Life Technologies). Cells were then washed in 1XPBS and mounted in Vectashield plus DAPI (Vector Laboratories). For combined immuno-RNA-FISH, immunofluorescence staining was performed first.

Rhesus FISH probes: rhesus *XIST* BAC (CH250-375L3), human *POLA1* BAC (RP11-11104L9), rhesus *ATRX* BAC (CH250-53K13), rhesus *FTX* BAC (CH250-372B10), rhesus *CHIC1* (CH250-234K8). BAC have been obtained from BACPAC resource center, CHORI. *JPX* probe consisted of six PCR products (see Supplementary Table1 for primer sequences).

Marmoset *XIST*, *JPX* and *ATRX* FISH probes consisted of PCR products (see Supplementary Table1 for primer sequences) that were concatenated by ligation before labelling.

All fluorescent microscopy images were taken on a fluorescence DMI-6000 inverted microscope with a motorized stage (Leica), equipped with a CCD Camera HQ2 (Roper Scientifics) and an HCX PL APO 100X oil objective (numerical aperture, 1.4, Leica) using the Metamorph software (version 7.04, Roper Scientifics). Approximately 40 optical z-sections were collected at 0.2 µm steps, at different wavelengths depending on the signal (DAPI [360 nm, 470 nm], FITC [470 nm, 525 nm], Cy3 [550 nm, 570 nm], and Cy5 [647 nm, 668 nm]). Stacks were processed using ImageJ and are represented as a 2D ‘maximum projection’ throughout the manuscript. Volume of the *XIST* clouds were measured with the Icy plateform for bioimage informatics (https://icy.bioimageanalysis.org/).

### RNA-seq data analysis

Publicly available RNA-Seq and scRNA-Seq data were retrieved from SRA and ENA, see Supplementary Table 2, and aligned using STAR on the rhesus macaque and marmoset reference genomes (rheMac10 and calJac4, respectively).

The macaque and marmoset orthologs of *XIST*, *FTX* and *JPX* were reconstructed using Scallop (0.10.4) with the default parameters^77^.

Read counts were quantified with htseq-count from htseq (0.13.5)^78^ using the following options: ––stranded reverse –a 10 –t exon –i gene_id –m intersection-nonempty. Custom annotation files for macaque and marmoset were created by adding *XIST*, *JPX* and *FTX* annotations to reference annotations files from ensembl.org, version 108. Reads marked as “no feature”, “ambiguous”, “too low aQual”, “not aligned” or “alignment not unique” were eliminated.

To determine the allelic expression, a custom version of the GATK pipeline for RNASeq short variant discovery was applied to RNA-Seq reads. The vcf were then filtered with stringent parameters to avoid false positive in SNP calling with VariantFiltration: – FS>60.0 –QD<2.0 –SOR>3.0 –MQ<40.0 –MQRankSum<-12.5 –MQPosRankSum<-8.0 and bcftools: –i ‘QUAL>=100 && INFO/DP>=10 && INFO/QD>=7’ –f PASS.

### Capture HiC

Tilling capture probes targeting a 3 Mb locus centred around the *XIST* gene in human (hg38, chrX:7241346275413462), rhesus macaque (rheMac10, chrX:69479389-70854356) and chimpanzee (panTro6, chrX:67854356-70854356) were designed and ordered through the Agilent Sure Design platform. *In situ* Capture HiC was performed as previously published^56^. Cells were crosslinked with 2% formaldehyde in PBS1X. 10^7^ cells were lysed on ice in 10 mM Tris–HCl pH8.0, 10 mM NaCl, 0,2% NP40, 1× complete protease inhibitor cocktail (Roche). Cells were de-crosslinked in 0,5% SDS in PBS1X at 37°C for 1h on a thermomixer under agitation. Nuclei were permeabilized using 1% Triton X-100 in H2O for 15 min at 37°C under agitation. 50 µl of nuclei suspension were collected for digestion control. Nuclei were then digested with DpnII (1:50) (NEB, 50 000 U/ml) for 4h at 37°C under agitation. Additional DpnII (1:50) was added and incubation was carried out at 37°C overnight on a rotating wheel. DpnII (1:50) was added again and nuclei were incubated for additional 4h at 37°C on a rotating wheel. DpnII was inactivated by heating at 65°C for 20 min and 50 µl were collected as digested non-ligated control. DpnII fragments were ligated with 240 Units of T4 ligase (Thermo Scientific 30 U/µl) overnight at 16°C on a rotating wheel. DNA was then purified by Proteinase K digestion (Eurobio) for 4h at 65°C, precipitated and incubated with RNaseA (Thermo Scientific). DNA quantities in sample and controls were determined using the Qbit broad range kit (Thermo Scientific) and 100 ng of each were loaded on an agarose gel as quality control of the digestion and ligation steps. The purified samples were sent to the EMBL Gene Core facility in Heidelberg, Germany, for capture, library preparation and sequencing (see^56^ for detailed protocol).

### Capture HiC data analysis

Sequencing reads from technical replicates were processed using the HiC-pro pipeline^79^, including, reads alignment on the human, macaque and chimpanzee reference genomes (hg38, rheMac10, panTro6 respectively with mapping quality>23) and valid interaction pairs detection. Raw contact matrices were built by binning in genomic windows of 2, 4, 6, 8 and 10 kb, using the allValidPairs2cooler.sh utility. Correlation analysis between the raw contact matrices of replicates was done using the hicCorrelate package from the HiCExplorer toolbox^80^. The 4 kb binning was retained based on the high correlation score and the contact matrices of both replicates were merged. The raw contact interaction frequencies of the captured region were normalized using the Iterative Correction approach^81^. Insulation score and domain boundaries were computed on the normalized contact matrices using the hicFindTads utility from the HiCExplorer suite with –minDepth 12000 –maxDepth 48 000 –step 8000. Viewpoints data were extracted using the chicViewpoint package from the HiCExplorer suite with default parameters.

### CUT&RUN

CUT&RUN was performed as previously described^82^. Briefly, 0.5 million cells per replicate were bound to 20 uL concanavalin A-coated beads (Bangs Laboratories) in Binding Buffer (20 mM HEPES, 10 mM KCl, 1 mM CaCl2 and 1 mM MnCl2). The beads were washed and resuspended in Dig-Wash Buffer (20 mM HEPES, 150 mM NaCl, 0.5 mM Spermidine, 0.05% Digitonin). The primary antibodies (1:50) were added to the bead slurry and rotated at RT for 1 hour. The beads were washed by Dig-Wash Buffer and pA-Mnase fusion protein (1:400, produced by the Institut Curie Recombinant Protein Platform, 0.785mg/mL) was added and rotated at RT for 15 min.

After two washes, the beads were resuspended in 150 µL Dig-Wash Buffer, and the MNase was activated with 2 mM CaCl2 and incubated for 30 min at 0 °C. MNase activity was terminated with 150 µL 2XSTOP (200 mM NaCl, 20 mM EDTA, 4 mM EGTA, 50 µg/ml RNase A and 40 µg/ml glycogen). Cleaved DNA fragments were released by incubating for 20 min at 37 °C, followed by centrifugation for 5 min at 16,000g at 4 °C and collection of the supernatant from the beads on a magnetic rack. The DNA was purified by phenol:chloroform and libraries were prepared using the TruSeq ChIP Library Preparation Kit from Illumina following the manufacturer’s protocol, and sequenced on a NovaSeq 6000 instrument (ICGex – NGS platform, Paris, France) to generate 2X100 paired-end reads. All the antibodies used in this study are listed in the Supplementary Table3.

### CUT&RUN data analysis

CUT&RUN reads were trimmed using trim_galore (0.6.5) (https://github.com/FelixKrueger/TrimGalore) with a minimum length of 50 bp. Reads were then mapped to the rhesus macaque genome (rheMac10) and mouse genome (mm10) using bowtie2 (2.4.4)^83^ with the following parameters: ––local ––very-sensitive-local ––no-unal ––no-mixed ––no-discordant ––phred33 –L 10 –X 700. Reads were then deduplicated using MarkDuplicates from picard (2.23.5) (http://broadinstitute.github.io/picard/) with the following options: –– CREATE_INDEX=true ––VALIDATION_STRINGENCY=SILENT –– REMOVE_DUPLICATES=true ––ASSUME_SORTED=true. Bam files were sorted, filtered (minimum mapping quality=10) and indexed with samtools (1.13)^84^. Reads from mouse embryonic fibroblasts (MEFs) were discarded using XenofilteR package from R (4.1.1)^85^. BigWig track files were generated with bamCoverage from deeptools (3.5.0) using the following parameters: ––normalizeUsing BPM ––binSize 20 –– smoothLength 40.

### ATAC-seq

ATAC-seq was performed as previously described^86^. Briefly, 50000 cells were resuspended in 50µL cold lysis buffer (10mM Tris-HCl; pH 7.4, 10mM NaCl, 3 mM MgCl2, 0.1 % Igepal CA-630) and centrifuged for 10 min at 500g at 4°C. The nuclei pellets were resuspended in 50 μl transposase reaction mix (25 µl 2X TD buffer, 2.5 µl transposase, and 22.5 µl H2O) and incubated at 37°C for 30 min in a thermomixer with 1000 RPM mixing. Reactions were cleaned up using the MinElute PCR purification kit (QIAGEN) and DNA was eluted in 10 µ elution buffer. Transposed DNA were pre-amplified for 5 cycles in 50 µl reaction mix (2.5 μl of 25 μM primer Ad1, 2.5 μl of 25 μM primer Ad2, 25 μl of 2X Master Mix, 10 µL H2O and 10 μl transposed elution) at the following cycling conditions: 72°C, 5 min; 98°C, 30 s; then 5 cycles of (98°C, 10 s; 63°C, 30 s; 72°C, 1 min). Then, 15 μl of qPCR amplification reaction (5 μl of pre-amplified sample; 0.5 μl of 25 μM primer Ad1, 0.5 μl of 25 μM primer Ad2, 5 μl of 2X NEBNext Master Mix, 0.24 μl of 25x SYBR Green in DMSO, and 3.76 μl of H2O) was carried out at the following cycling conditions: 98°C, 30 s; then 20 cycles of (98°C, 10 s; 63°C, 30 s; 72°C, 1 min). The required number of additional cycles for each sample was determined as described previously^86^. After the final amplification, double-sided bead purification was performed with AMPure XP beads. Final ATAC-seq libraries were eluted in 20 μl nuclease-free H2O from the beads and were sequenced on a NovaSeq 6000 instrument (Novogene, UK) to generate 2X150 paired-end reads.

ATAC-seq raw data were analyzed using the atacseq pipeline (1.2.1) (https://github.com/nf-core/atacseq/) from nf-core^87^ using the default parameters. Briefly, reads were aligned to the macaque genome (rheMac10) using BWA (0.7.17). Duplicates were marked by picard and reads mapping to mitochondrial DNA and blacklisted regions were removed. BigWig files were generated using deeptools as described above.

### Peak calling, chromatin state and binding motif annotation

Peak calling was performed on the bam files generated from the CUT&RUN, ChIP-Seq, ATAC-Seq and DNAse-Seq listed in Supplementary Table 4 using MACS2 with – g 2.7e9. Only peaks with FDR<0.05 were retained.

Chromatin states were called by processing H3K4me3, H3K4me1, H3K27Ac CUT&RUN and chromatin accessibility (ATAC-Seq, rhESC; DNAse-Seq, hESC) with chromHMM^61^. Loci of high H3K4me3/chromatin accessibility were annotated as “promoter”, loci with high H3K4me1/H3K27Ac/chromatin accessibility were annotated as “active enhancer”, loci with high H3K27ac and low H3K4me1 were annotated as “poised enhancer” as previously reported^88^.

CTCF motifs were annotated using the FIMO tool from the MEME suite (5.5.1)^89^ with the CTCF matrix profile (MA0139.1) downloaded from JASPAR. Only motif with FDR<0.001 were kept.

### CRISPR-Cas9 deletion and CRISPR-inhibition

SgRNAs for both CRISPR-Cas9 and CRISPRi experiments were designed using the web-based tool CRISPOR (http://crispor.tefor.net/) and only the sgRNAs with the least probability of off-target were selected. SgRNAs designed for CRISPR-Cas9 deletions were cloned into the pSpCas9(BB)-2A-GFP vector (Addgene #48138) or into the pSpCas9(BB)-2A-mCherry vector as previously described^41^. SgRNAs designed for CRISPRi experiments targeting the *JPX* promoter were cloned into the PB_rtTA_BsmBI vector (Addgene, #126028) and sgRNAs targeting Enh5 were cloned into a multiexpression sgRNA vector, a gift from the Schulz lab^90^. All sgRNA sequences can be found in Supplementary Table1 for rhesus macaque and Table2 for marmoset.

For transfection, cells were dissociated with Accutase (Stemcell Technologies) to obtained single-cell suspension. Approximatively 10^6^ cells were nucleofected using the Amaxa 4D-Nucleofector^TM^ system (Lonza). For CRISPRi-based strategies, the PB_rtTA_BsmBI vector, PB_tre_dCas9_KRAB (Addgene, #126030), and piggyBac transposase^91^ were delivered to the cells in a 1:1:2 ratio. For CRISPRi-based strategies, rhESC positive clones were selected with G418 (300 µg/mL) and hygromycin (50 µg/mL) and positive cjESC were selected with G418 (200 µg/mL) and hygromycin (50 µg/mL). The number of random insertions in the genome was verified by qPCR and clones with the lowest insertion number (generally between 2 and 3 insertions of each vector per cell line) were used for further experiments. For induction, cells were treated with 1µg/mL doxycycline for 5 to 10 days. For deletion-based strategies, clones were screened by PCR (see supplementary Table1 for primer sequences) then phased by Sanger sequencing.

### Total RNA extraction and RTqPCR

Total RNA was purified using RNeasy mini kit (QIAGEN) following the manufacturer’s recommendation and quantified on a nanodrop 2000. 1 μg of RNA were treated with DNaseI (TURBO™ DNase, Thermo Fisher Scientific) for 30 min at 37°C and reverse transcribed with the reverse transcriptase Superscript IV kit (Thermo Fisher Scientific) following the manufacturer’s recommendation. cDNAs were diluted 2.5 times in water and RNA expression level was assessed by real time quantitative PCR using the Power SYBR Green Master Mix (Thermo Fisher Scientific) on a ViiA-7 Real-Time PCR thermocycler (Applied Biosystems). RNA levels were normalized against Beta-actin reference gene following the 2-ΔCt method. For qPCR primer sequences see Supplementary Table 1 and Table 2.

### ChIP and qPCR

For chromatin preparation, a minimum of 1 million ESCs were fixed with 1% Formaldehyde (Electron Microscopy Sciences) for 10 min on a shaker at room temperature and quenched with Glycine 0.125M for 5 min at room temperature. Cells were then washed with PBS1X and cell pellets were flash-frozen and stored at –80°C. Samples were lysed in (50 mM Tris-HCl pH7; 10 mM EDTA; 1% SDS) supplemented with protease inhibitors (Thermo Fisher Scientific) and sonicated using a Bioruptor Pico (Diagenode) and 15 cycles (30 sec OFF/30 sec ON). For ChIP, samples were incubated overnight with 5 μg of H3K27ac (Active Motif 39034) or with 5 μg of H3K9me3 (Active Motif 39062) per 20 μl of protein A/G MagnaBeads (Merck Millipore). Beads were then washed once with low salt wash buffer (0.1% SDS; 1% Triton X100; 2 mM EDTA; 20 mM Tris-HCl pH8; 150 mM NaCl), once with high salt wash buffer (0.1% SDS; 1% Triton X100; 2 mM EDTA; 20 mM Tris-HCl pH8; 500 mM NaCl) and once with LiCl wash buffer (0.25 M LiCl; 1% NP-40; 1% Sodium Deoxycholate; 1 mM EDTA; 10 mM Tris-HCl pH8) and treated with 1 mg/mL of proteinase K for 45 min at 50°C under agitation. DNA was purified using GeneJET Gel Extraction kit (Thermo Fisher). For qPCR primer sequences see Supplementary Table 1 and Table 2.

### Sequence analyses

To date the HERVK insertion within the primate *XIC,* a custom BLAST database spanning CHIC1-XIST orthologous intervals was constructed from NCBI retrieved sequences. HERVK consensus sequence from Dfam was blasted against it. To date the HERVK invasion of the primate genomes, the consensus was blasted against the genome of the selected primates. All phylogenetic trees were created with www.timetree.org. All species silhouettes were taken from https://www.phylopic.org/.

The phyloP scores for human and macaque were generated using the reference-free multi-way alignment of 241 mammalian species^2^. The annotations of constrained and not constrained CTCF motifs in hESCs were generated using the resource referencing the natural selection over every human CTCF sites^16^. hESC constrained enhancer list was built by overlapping the active enhancers from the present analysis with intergenic regions under selective constraint in humans, that is the zooUCE, UNICORNS and RoCCs loci identified by the Zoonomia consortium^2^. TSS coordinates for human and macaque genes were downloaded as bed files from CAGE-Seq peaks generated by the FANTOM5 consortium https://fantom.gsc.riken.jp/5/datafiles/phase2.6/extra/CAGE_peaks. CAGE-Seq peaks were overlapped with gene annotation from ensembl.org to annotate the biotype, i.e., protein-coding or lncRNA, of each TSS. The sum of the phyloP scores spanning each feature was then computed for chromosome 7, 8 and X.

### Data and code availability

All the data generated in this study has been deposited on GEO under the accession number GSE248841. All the publicly available datasets used in this study are listed in Supplementary Table 4. The code supporting the analysis is available upon request to the corresponding authors.

## Acknowledgements

We thank lab members for critical evaluation of the work leading to this publication. We thank Gregory Andrews from UMass Chan Medical School for sharing the annotation of human constraint CTCF sites. We thank Dr. Magali Hennion for bioinformatics support provided at the Bioinformatics and Biostatistics Core Facility (BiBs), Paris Epigenetics and Cell Fate Center. We thank the (EPI)^2^ Imaging platform – UMR7216 Epigenetic and Cell Fate Centre – for access to instruments and technical advices. We thank the Functional Epigenomics (EpiG) platform hosted at the UMR7216 Epigenetic and Cell Fate, for technical advices and access to instruments. We acknowledge the ImagoSeine core facility of the Institut Jacques Monod, member of the France BioImaging infrastructure (ANR-10-INBS-04) and GIS-IBiSA, in particular, we thank the Imagoseine cytometry platform and especially Nicolas Valentin for technical assistance and advices. We thank the ICGex NGS platform of the Institut Curie supported by the grants ANR-10-EQPX-03 (Equipex) and ANR10-INBS-09-08 (France Génomique Consortium) from the Agence Nationale de la Recherche (“Investissements d’Avenir” program), by the ITMO-Cancer Aviesan (Plan Cancer III) and by the SiRIC-Curie program (SiRIC Grant INCa-DGOS-465 and INCa-DGOSInserm_12554) for the high-throughput sequencing. We thank the Bioinformatics platform of the Institut Curie for data management, quality control and primary analysis, in particular Dr. Nicolas Servant, Head of the platform, for advices on capture-HiC data analyses. This project was funded by Agence Nationale pour la Recherche [ANR-14-CE10-0017 to C.R. and to P.S.]; European Research Council (ERC) under the European Union’s Horizon 2020 research and innovation program [101020423]; LabEx ‘Who Am I?’ [ANR-11-LABX-0071]; Université Paris-Cité IdEx [ANR-18IDEX-0001] funded by the French Government through its ‘Investments for the Future’ program. E.C. was supported by fellowships from the French Ministry of Education and Research and from the French Medical Research Foundation (FRM).

