## Supplementary figures and images for "Extensive remodelling of *XIST* regulatory networks during primate evolution"

### Supplementary figures

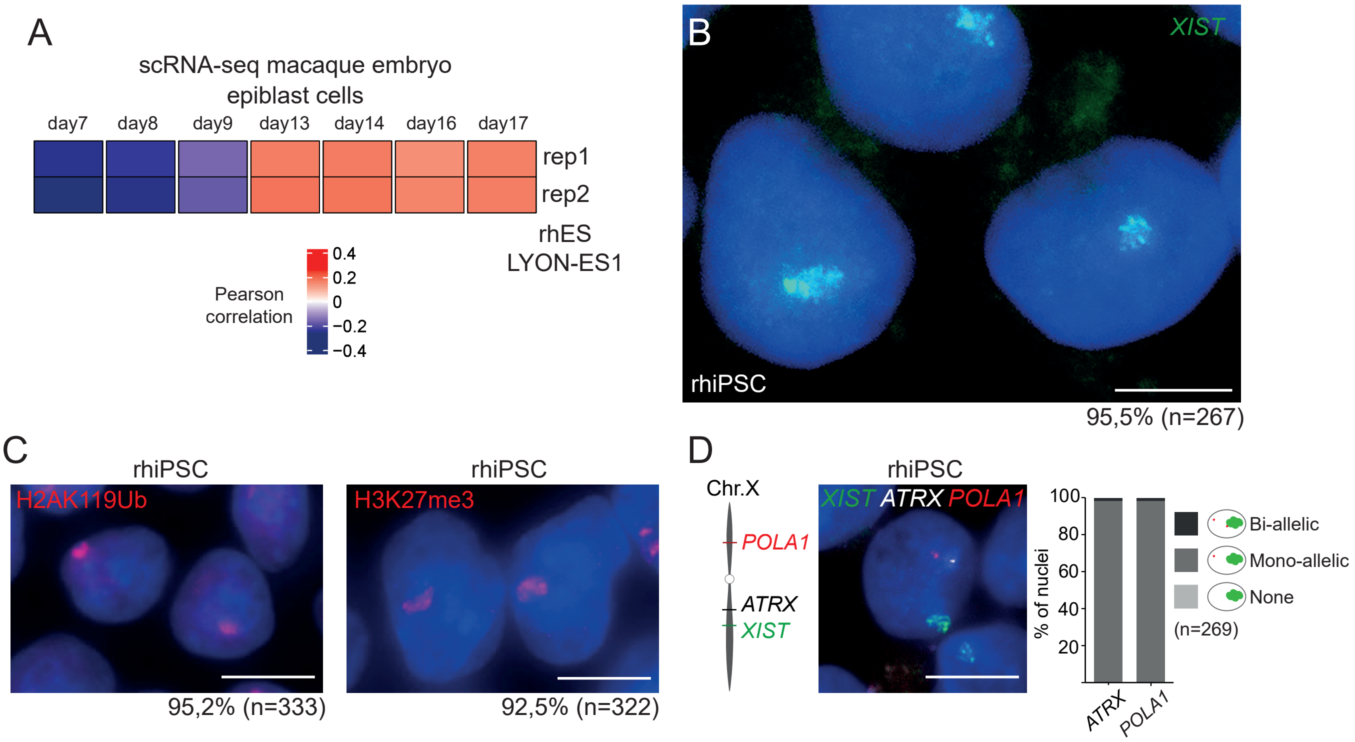

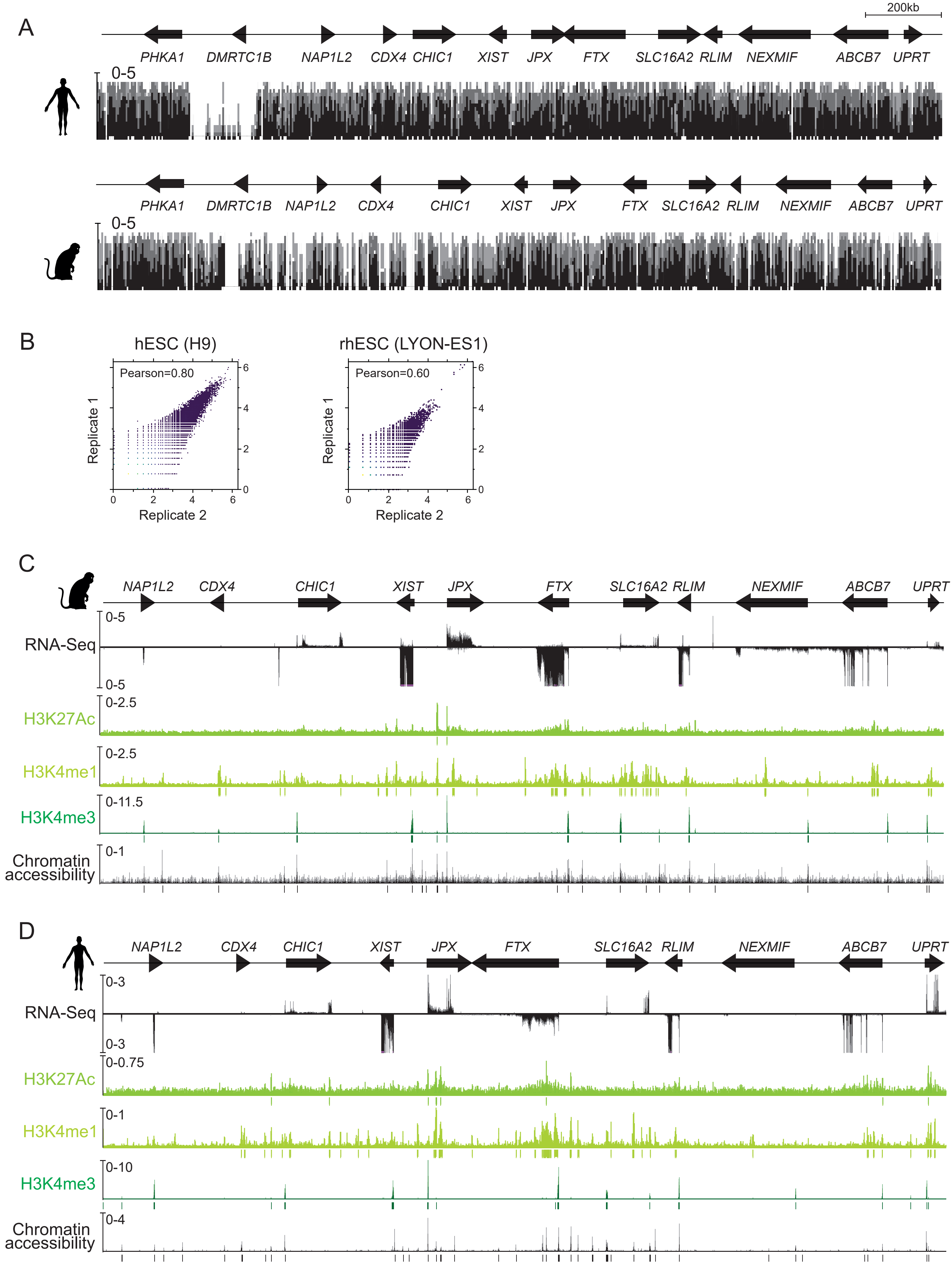

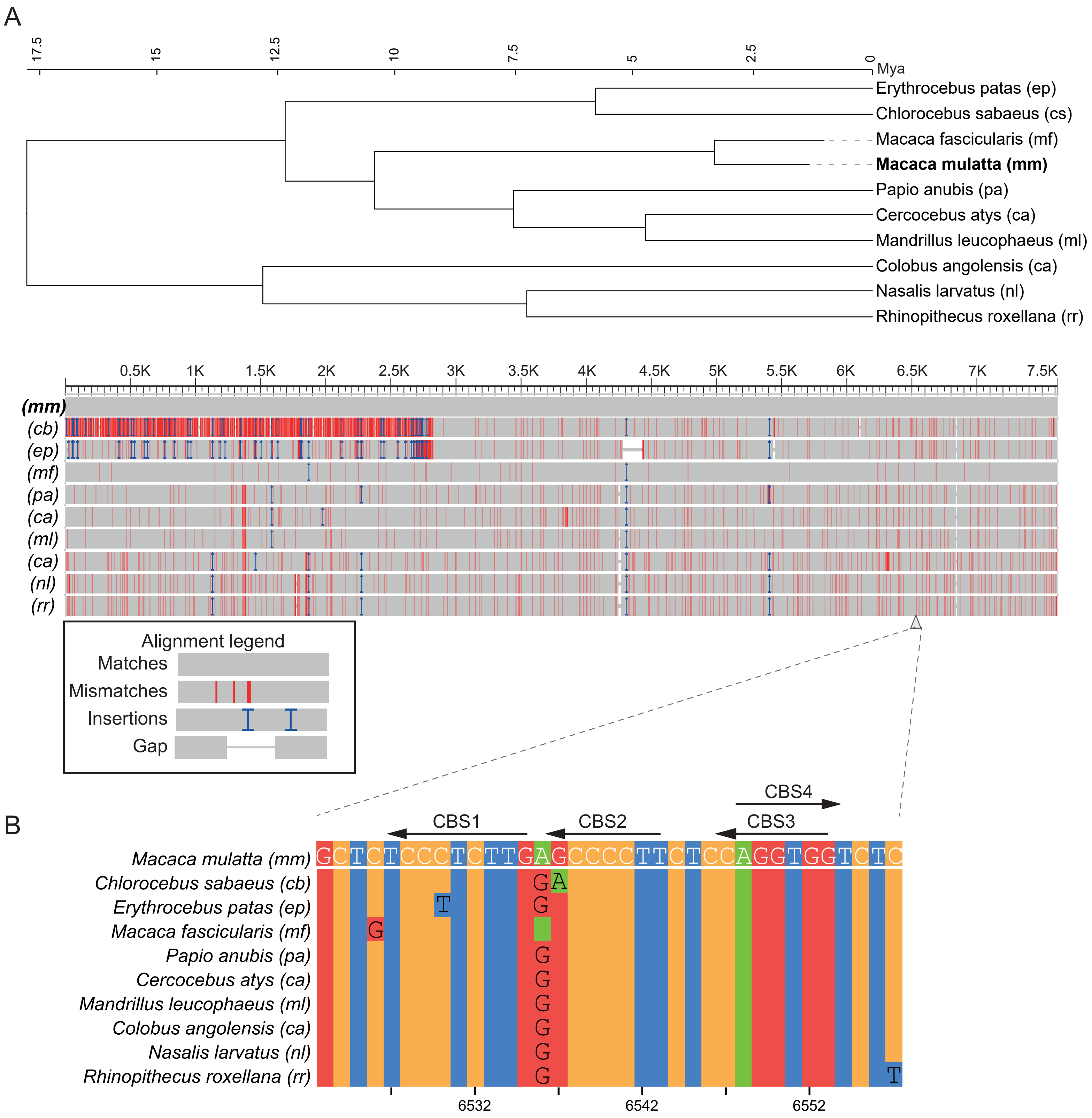

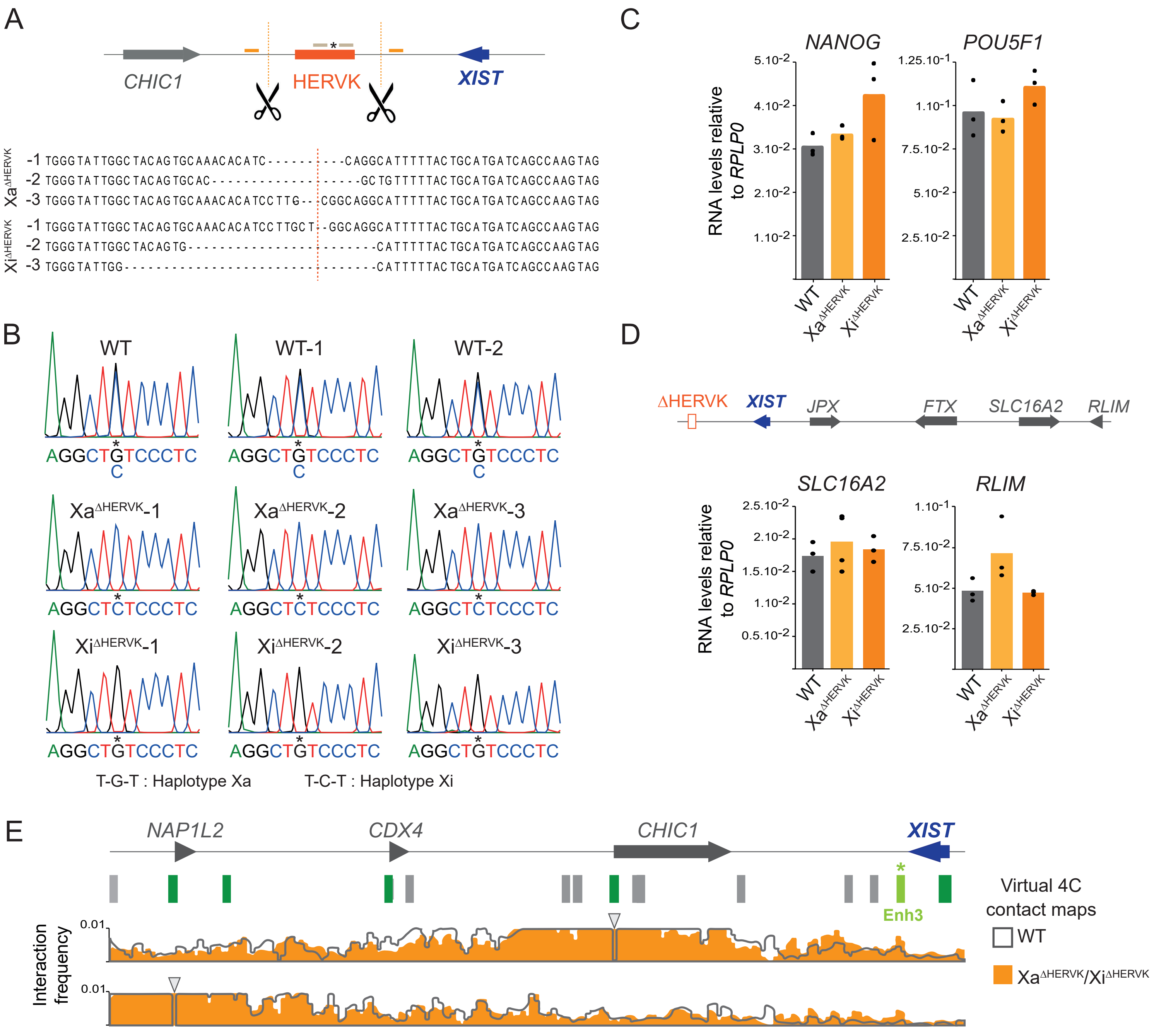

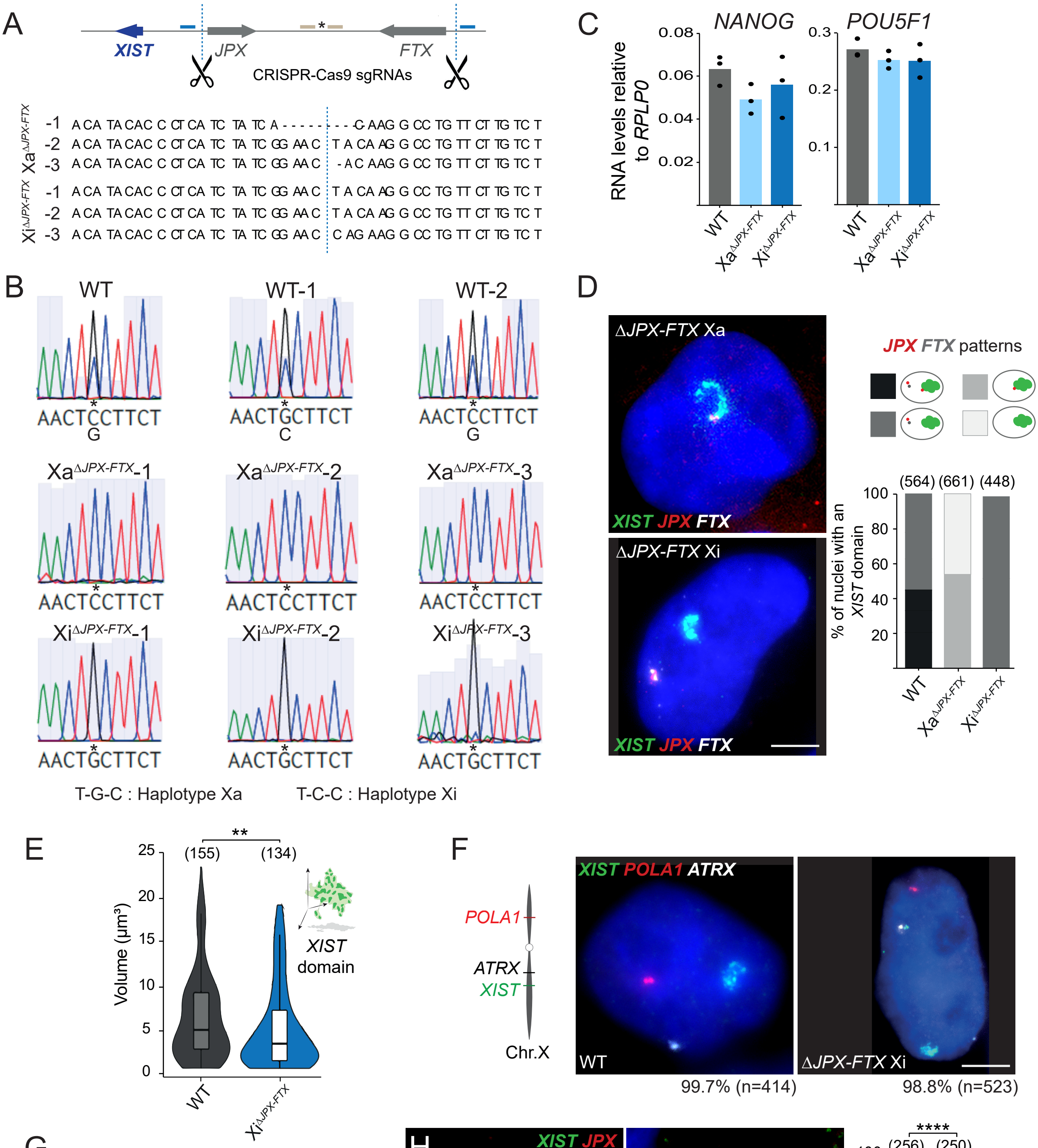


G

| Clone | rTA copy number | dCas9 copy number |
|---|---|---|
| 1 | 2 | 3 |
| 2 | 2 | 2 |
| 3 | 2 | 2 |

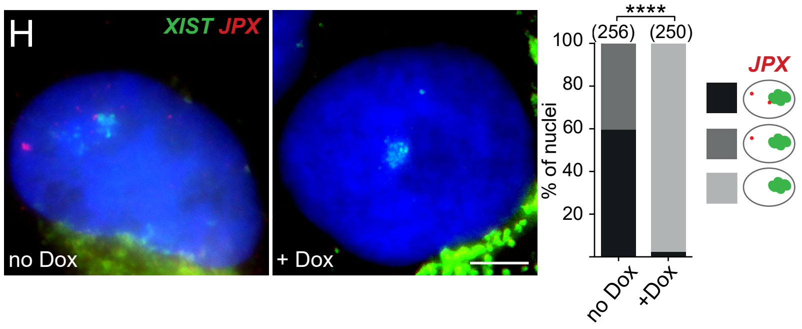


Supp 5

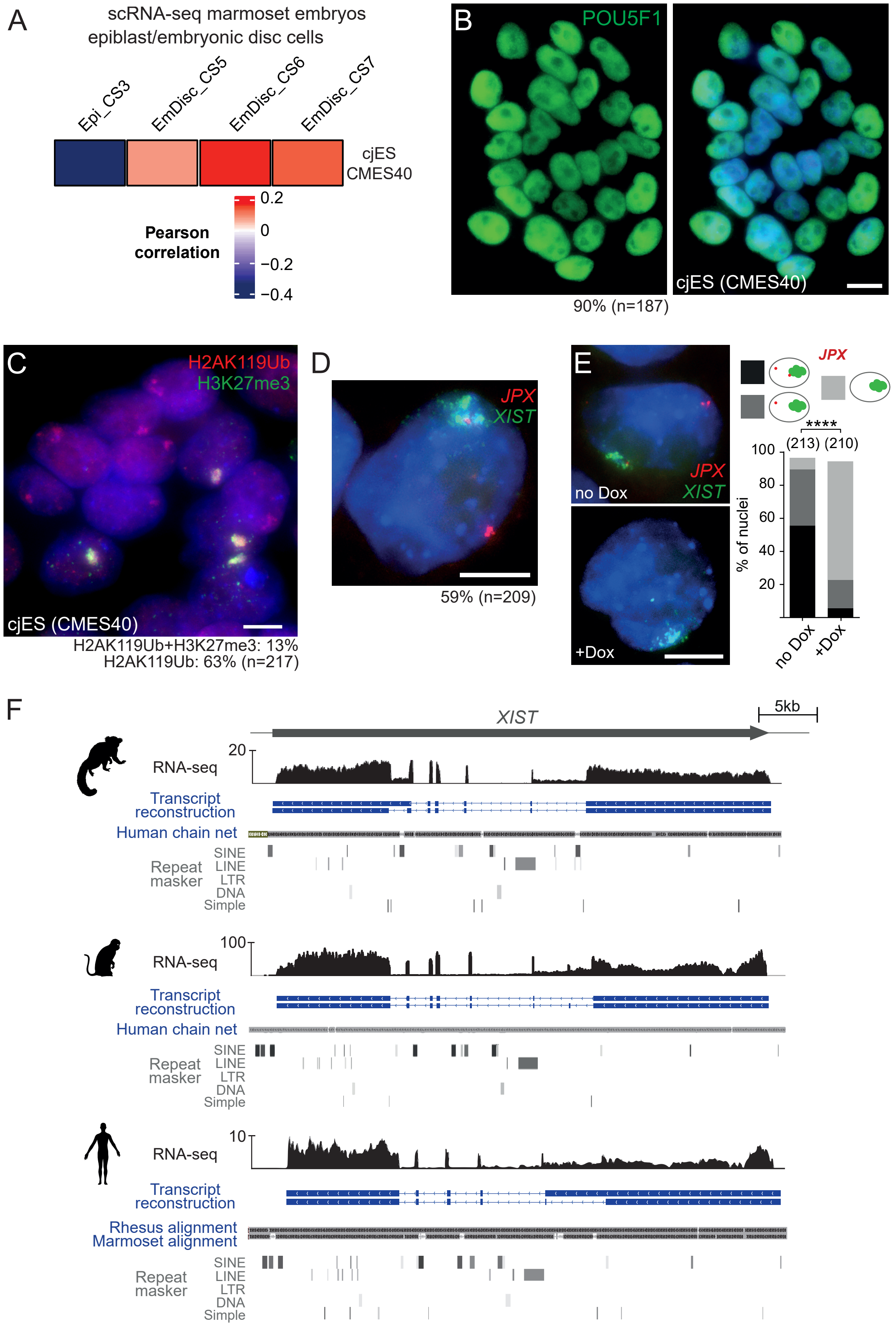

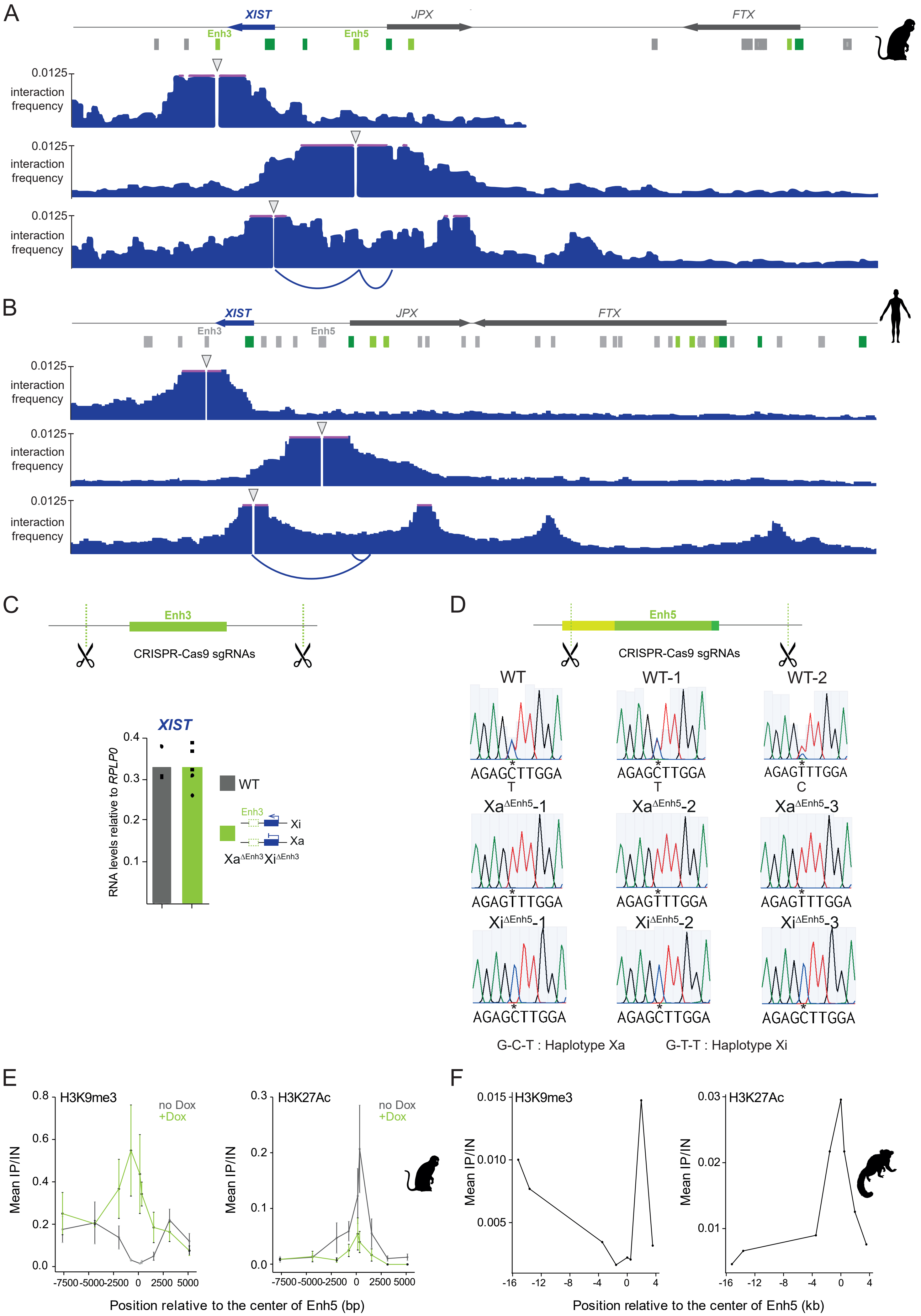

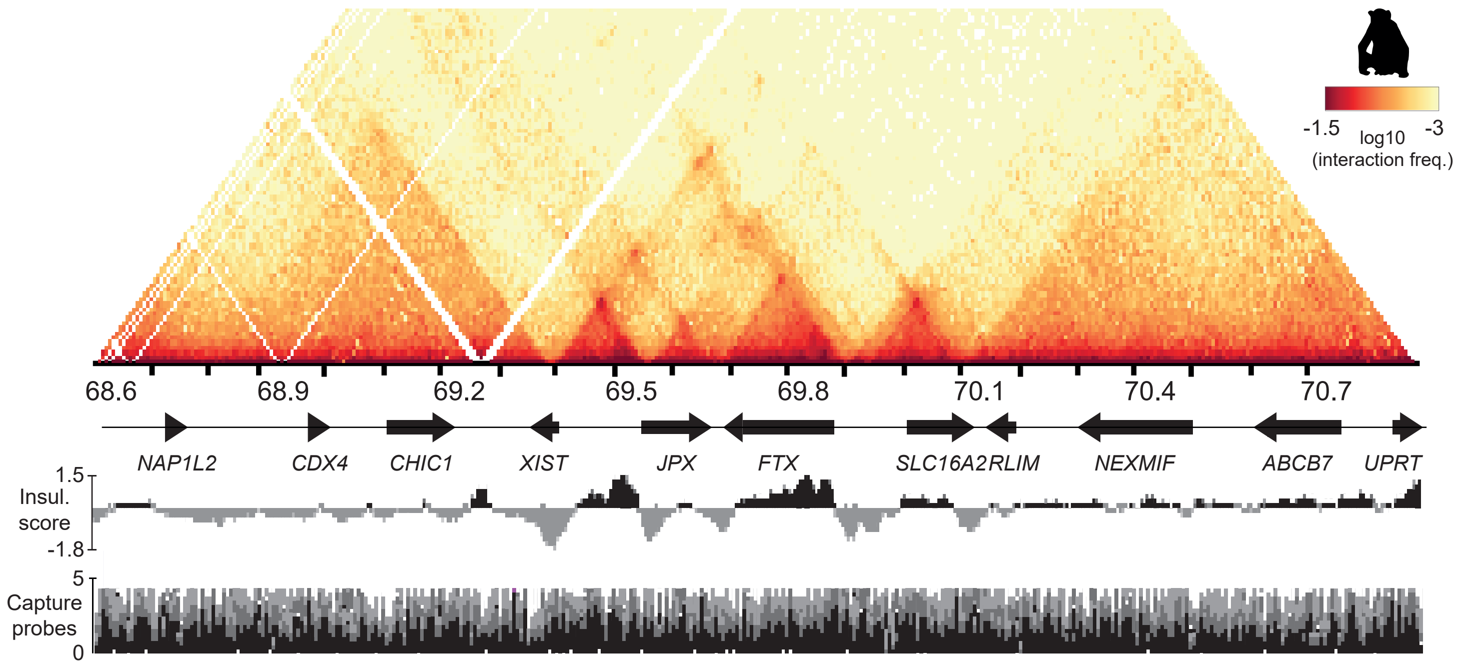
